# Microbial ecology of sulfur biogeochemical cycling at a mesothermic hot spring atop Northern Himalayas, India

**DOI:** 10.1101/2021.12.02.470874

**Authors:** Shekhar Nagar, Chandni Talwar, Mikael Motelica-Heino, Hans-Hermann Richnow, Mallikarjun Shakarad, Rup Lal, Ram Krishan Negi

**Author notes:** Corresponding Author, Department of Zoology, University of Delhi, Delhi 110007, India.

## Abstract

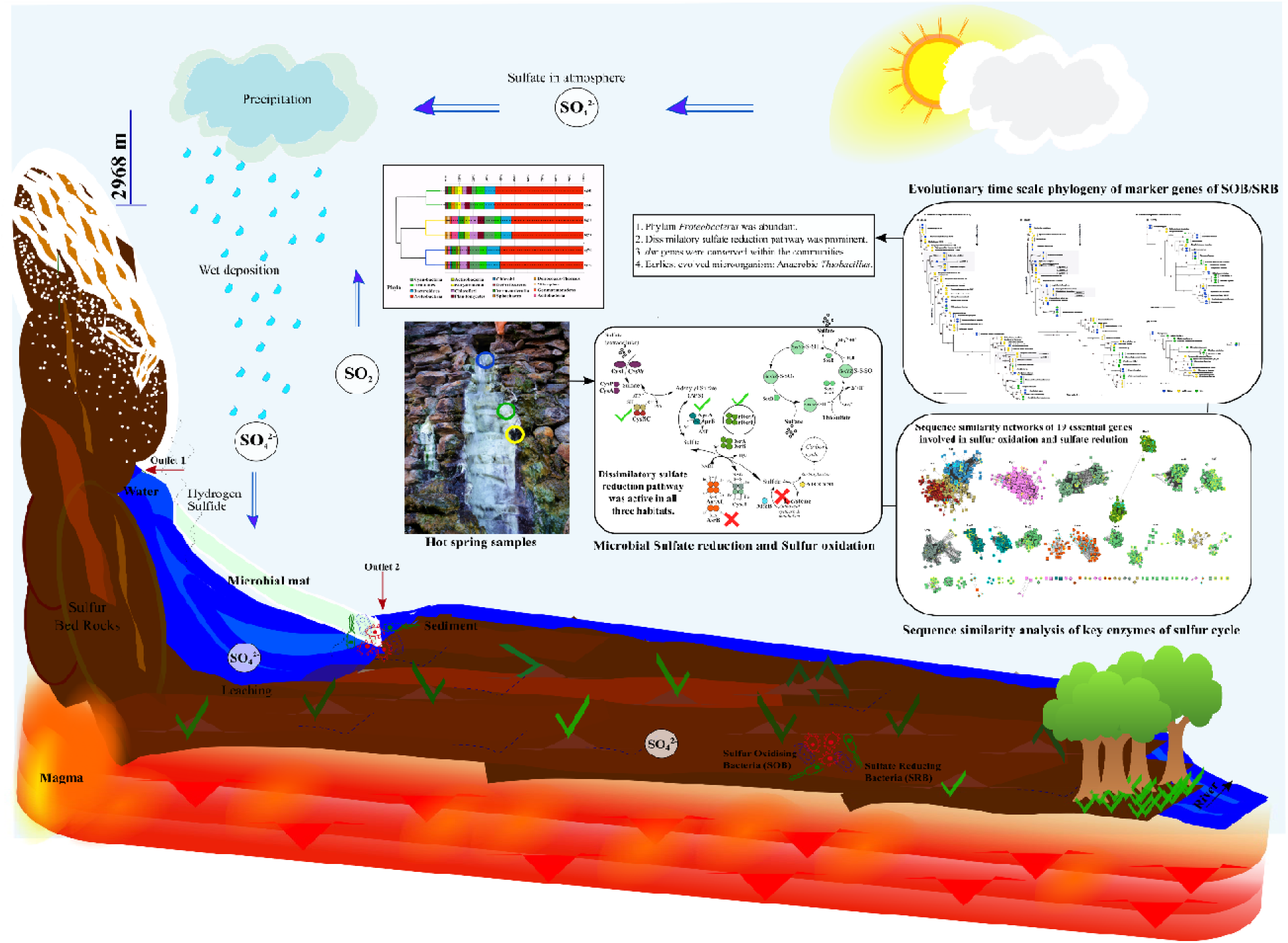
Graphical Abstract.

Sulfur Related Prokaryotes (SRP) residing in hot spring present good opportunity for exploring the limitless possibilities of integral ecosystem processes. Metagenomic analysis further expand the phylogenetic breadth of these extraordinary sulfur metabolizing microorganisms, as well a their complex metabolic networks and syntrophic interactions in environmental biosystems. Through this study, we explored and expanded the microbial genetic repertoire with focus on sulfur cycling genes through metagenomic analysis of sulfur (S) contaminated hot spring, located at the Northern Himalayas. The analysis revealed rich diversity of microbial consortia with established roles in S cycling such as *Pseudomonas*, *Thioalkalivibrio*, *Desulfovibrio* and *Desulfobulbaceae* (*Proteobacteria*). The major gene families inferred to be abundant across microbial mat, sediment and water were assigned to *Proteobacteria* as reflected from the RPKs (reads per kilobase) categorized into translation and ribosomal structure and biogenesis. Analysis of sequence similarity showed conserved pattern of both *dsrAB* genes (n=178) retrieved from all metagenomes while other sulfur disproportionation proteins were diverged due to different structural and chemical substrates. The diversity of sulfur oxidizing bacteria (SOB) and sulfate reducing bacteria (SRB) with conserved (r)*dsrAB* suggests for it to be an important adaptation for microbial fitness at this site. Here, we confirm that (i) SRBs belongs to *δ-Proteobacteria* occurring independent LGT of *dsr* genes to different and few novel lineages (ii) also, the oxidative and reductive *dsr* evolutionary time scale phylogeny, proved that the earliest (not first) *dsrAB* proteins belong to anaerobic *Thiobacillus* with other (*rdsr*) oxidizers. Further, the structural prediction of unassigned DsrAB proteins confirmed their relatedness with species of *Desulfovibrio* (TM score= 0.86; 0.98; 0.96) and *Archaeoglobus fulgidus* (TM score= 0.97; 0.98). We proposed that the genetic repertoire might provide the basis of studying time scale evolution and horizontal gene transfer of these genes in biogeochemical S cycling and the complementary genes could be implemented in biotechnology and bioremediation applications.

## Introduction

The untapped sulfur compounds oxidizing microorganisms (SOM) and sulfur compounds reducing microorganisms (SRM) microbial communities residing in extreme and contaminated environmental conditions such as hot water, sulfide contaminated springs offer an intriguing opportunity to explore the unique microbial diversity with uncovered metabolic potential (Ghilamicael et al., 2017). The investigations of such microbiota began with focus on identifying and culturing novel thermostable biocatalysts with huge biotechnological applications (Inskeep et al., 2010, Li et al., 2014, Ayangbenro and Babalola., 2017). Over the decades, metagenomics studies have revealed the crucial role of abiotic ecology that guides the microbiota at such sites (Lozupone et al., 2007, Inskeep et al., 2013). Evidences of such environmentally adapted microbiota are vast in terms of diversity and its correlation with temperature (Blank et al., 2002; Miller et al., 2009; Reigstad et al., 2010; Ghosh et al., 2012; Cole et al., 2013; Chan et al., 2015). However, little progress has been made in exploring the correlation between microbiome and geochemistry of hot spring systems particularly that possess mesothermic hot waters with neutral pH and elemental sulfur and sulfate richness (Roy et al., 2020; Ghosh et al., 2012). Moreover, the survival of microbes in these niches is often supported by community dynamics and interactions. Studies of such ecosystems may provide insights into the microbial evolution of specific pathways for microbial biogeochemical cycling of minerals. However, with over 400 thermal hot water springs located in India, less than 15% have been explored for biogeochemical and taxonomical classification using genomics and metagenomics approaches (Cinti et al., 2009; Saxena et al., 2017). Sulfur springs provide harsh physiochemical conditions to sustain the growth of only meso and hyper-thermophilic microbes which includes sulfur oxidizers, sulfate reducers (Chan et al., 2015; Gonsior et al., 2018). The survival could also be achieved with “microorganism adaptation” by several resistance mechanism such as activity of bioprecipitation, biosorption, extracellular sequestration and/or chelation (Haferburg and Kothe 2007). During these changes, the exchange of genetic material by means of horizontal gene transfer (HGT) is prevalent and necessary for the adaptation of microbes through acquisition of novel genes.

Khirganga, the mesothermal sulfur spring in Northern Himalayas discharging waters rich in sulfate, chlorine, sodium and magnesium ions has remained uncharted so far (Shirkot and Verma 2015; Poddar and Das, 2018). High levels of sulfides in the environment accounts for the milky appearance of the hot spring water with white microbial mats predicted to be formed from sulfide reduction by the sulfur related prokaryotes (SRP) enriched at this site (Sharma et al., 2004; Dong et al., 2019). The microbial disproportionation one of the oldest (3.5 billion years ago; Finster 2011) biological processes on Earth producing sulfide, sulfite and sulfate compounds establishes a complex network of pathways in the biogeochemical sulfur cycle. Heretofore, it is the very foremost metagenomic investigation of microbial communities in Khirganga (avg. atmospheric temperature 6.9 ± 0.3) focused on exploring the microbial biogeochemical sulfur cycling with a complex of disproportionation of elemental sulfur conforming intermediary compounds. The current study was carried out via microbial mats, sediments and hot spring water samples in hot spring to decipher the stabilized and diversified genes involved in sulfur cycle intermediary process in sulfate-reducing bacteria and anoxygenic, photolithotrophic and chemolithotrophic sulfur-oxidizing bacteria (Dahl & Truper, 1994; Hipp et al., 1997). The work expands the genetic and evolutionary information for sulfur cycling genes and evaluates the biodiversity and applications for screening of the novel thermostable enzymes from microorganisms. Further, understanding these adaptations vis-à-vis the physiological properties and metabolic processes in these springs could be monitored as the engineered SRP consortia could develop into an effective tool in optimizing degradation of sewage waste in industrial processes (Ayangbenro et al., 2018). Also, the sulfate-reducing bacteria (SRB) implied to treat various environment contaminants including metals (Zhang et al., 2016, Mothe et al. 2016), metalloids (Bataglia-brunet et al., 2012, Sahinkaya et al. 2015), various non-methane hydrocarbons (Callaghan et al., 2012), alicyclic hydrocarbons (Jaekel et al., 2015), nitroaromatic compounds (Boopathy 2014, Mulla et al., 2014) and aromatic hydrocarbons (Stasik et al., 2015, Meckenstock et al., 2016, Kamarisima et al., 2019).

## Material and Methods

### Sample collection, Physicochemical analysis and Helium Ion Microscopy

Samples of water (5L), sediment (250 g) and microbial mat deposits (250 g) were collected from Khirganga hot water spring (31°59*’*34*"* N, 77°30*’*35*"* E) in February, 2017. Sampling was performed in two replicates for each habitat from two closely located primary thermal outlets (31°99*’*18*"* N, 77°50*’*96*"* E) and secondary outlets (31°99*’*19*"* N, 77°50*’*96*"* E). The surface temperature and pH of each habitat were recorded on site.

Priorly, water samples were filtrated to 0.1 µm, sediment and microbial mats were digested in pure nitric acid prior to chemical analysis. All samples were subjected to physicochemical analysis for major elements ncentrations of major cations Na^+^, K^+^, Mg^2+^ and Ca^2+^) and anions (SO_4_ ^2-^, and Cl^-^ ) were analyzed by ionic chromatography (Dionex ICS-2000, Sunnyvale, CA) using the columns CS16A for measuring cations and AS17 for anions. Elemental analysis of minor and trace elements through inductively coupled plasma mass spectrometry (ICP-MS) Agilent ICP-MS 7900 with Ultra High Matrix Introduction (UHMI). Samples of sediments and microbial mats were desiccated overnight followed by ethanolic dehydration and microstructure was studied on scanning electron microscope at the Centre for Chemical Microscopy (ProVIS). Images were captured using high efficiency detector.

### Metagenomic DNA Extraction, Sequencing and Assembly

Total community DNA from 5 liters of filtered water (0.45µm) and 0.25g sediment samples were extracted using PowerMax Soil DNA isolation kit (MoBio Laboratories Inc., Carlsbad, CA, USA) following the manufacturer’s instructions. For extraction of total DNA from microbial mats, 0.25g samples were processed following a method described by Varin *et al.,* 2010. Sequencing was performed at Beijing Genome Institute (BGI), Hongkong, China using Illumina Hiseq 2500 platform. Paired end libraries of read length 100bp were generated with insert size of 350bp. The raw sequences were quality filtered using SolexaQA (Cox et al., 2010), and the low-quality sequences <Q_20_ quality cut-off and artificially duplicated reads (ARDs) were castoff using Illumina-Utils (Eren et al, 2013) and duplicate read inferred sequencing error estimation (DRISEE) (Gomez-Alvarez et al., 2009), respectively. Further, the assembly was integrated in IDBA-UD (Peng *et al.,* 2012) with insertion length 50 bp, min. *k-mer*: 31, max. *k-mer*: 93 (61 for water) using seed *k-mer* size for alignment 30 bp and min. size of contig as 200 bp while allowing min. multiplicity for filtering *k-mer* while building the graph.

### Taxonomic and Functional Assignments

Alpha diversity within each sample was estimated as abundance-weighted average of annotated species from source databases built in MG-RAST v3.0 (Meyer *et al.,* 2008) and expressed as Shannon diversity index transformed based on rarefaction curve using the following formula:

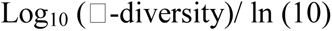

Diversity at phylum level was inferred from MG-RAST (maximum e-value, 1× 10^5^ and minimum percentage identity cutoff, 60%). A paired-sample t-test was applied on the phylum determined in any habitat pair to estimate significant similarities based on taxonomic mean abundance using SPSS (SPSS Inc., version 20.0, IBM). Microbial genera were deciphered based on clade-specific markers to identify taxonomy up to species level using MetaPhlAn v2.0 (Truong *et al.,* 2015) and a heatmap was constructed using Bray-curtis dissimilarity with supporting dendrograms for both species and samples. We used HUMAnN2 (Franzosa *et al.,* 2018) to perform phylum-resolved functional profiling of the communities that maps contigs onto the pangenomes of the known species of the community and quantifies the pathways and uniref90 gene families database (Suzek *et al.,* 2015). Later, these UniRef90 families were regrouped as COG annotations based on eggNOG (Huerta-Cepas *et al.,* 2017). Open Reading Frames (ORFs) of assembled metagenome were predicted using Prodigal v2.6.1 (Hyatt *et al.,* 2010) and annotated at hierarchy levels *viz.* subsystems, protein families and individual enzymes using Prokka v1.12 (Seemann, 2014). The amino acid sequences were mapped against KEGG database (Kanehisa *et al.,* 2004) and top 50 metabolic pathways in all of the six samples were compared through heatmap constructed using package ‘*pheatmap*’ (Kolde and Kolde, 2015) and ‘*ggplot2*’ (Wickham, 2009) in R (R Development Core team). Identification of S substrates disproportionation genes was performed by mapping all predicted ORFs on the HMM databases obtained from TIGRfam v10 (Haft *et al*., 2012) and Pfam (Finn et al., 2014) using hmmscan v3.1b2 (Eddy, 2011). Abundance of each enzyme was plotted as number of copies annotated within each sample. The sequences with >150 amino acids were queried against the NCBI Microbial proteins from RefSeq *nr* database (4^th^ April, 2020) using Blastp (Altschul et al., 1990) to identify the sequences producing significant alignments for taxonomic confirmation.

### Analysis of diversity of sulfate reduction proteins

For sequence similarity networks (SSN), amino acid sequences of sulfide oxidation and sulfate reduction proteins annotated in all six samples annotated by KEGG Ids were implied over an empirical measurement of diversity. For this, an all-vs-all BLAST was performed to define the similarities/variations between sequence pairs of diversifying sulfate reduction proteins. A user defined threshold was optimized according to the alignment score and maximum length of BLAST results in diversifying and stabilized protein sequences. Clustering was performed using CD-HIT (Li & Godzik, 2006) on the scores of BLASTp pairwise alignments at a threshold value (e-value of 1e-30). The networks were visualized in Cytoscape v3.7.1 (Shannon *et al.,* 2003). The average number of degree and neighbours for a protein sequence or a node was calculated under:

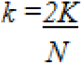

Where K is denoted with number of edges and N is denoted with total number of nodes. Also, to determine the divergence/similarity among nodes or protein sequences was calculated under:

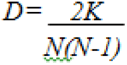

Attributes of node degree distribution, average clustering coefficient, average neighborhood connectivity and closeness centrality were studied through power law fits to determine their correlation with number of neighbors. Sulfur oxidizing bacteria (SOB) and sulfate reducing bacteria (SRB) were identified for the sequences that could be classified up to genus level to study the distribution of S substrates oxidation and reduction genes in the different clusters.

### Sequence alignment, phylogeny and structure prediction of putative unidentified Dsr and Asr enzymes

To elucidate the diversity of key sulphite reductases, the DsrA/B and AsrA/B protein sequences were individually aligned using MUSCLE v3.8.31 (Edgar 2004) and clustered using UPGMB. All the alignments were end trimmed manually and maximum likelihood (ML) phylogeny was inferred with 500 bootstrap resampling using RAxML v8.0.26 (Stamatakis 2014). For this, we used standalone version of RAxML which was called as follows:

raxmlHPC-PTHREADS -s input -N 500 -n result -f a -p 12345 -x 12345 -m PROTGAMMAGTR.

The resulting phylogenies were also confirmed using most complex general time-reversible model (GTR; Tavare, 1986) with PhyloBayes v1.7b using CIPRES Science Gateway v 3.3 (Lartillot et al., 2009) that incorporates different rates for every change and different nucleotide frequencies.

For proteins showing similarity with those from uncultured bacteria, we determined the structures using I-TASSER suite (Yang and Zhang, 2015). These predicted structures were then aligned onto their top structural analogs and C-scores, TM-scores and RMSD were computed and ligand binding sites with conserved residues were identified. The TM-score is to compare two models based on their given residue equivalency (i.e., based on the residue index in the PDB file). It is usually NOT applied to compare two proteins of different sequences. The TM-score predicted from structural alignment of two proteins while comparing them based on residue equivalency such that a score of 0.6 and above denote the two proteins to be fairly aligned (Yang and Zhang, 2015). The TM-align will first find the best equivalent residues of two proteins based on the structure similarity and then output a TM-score.

## Results and Discussion

### Description of sampling site and microscopic analysis of samples

The Khirganga is a natural hot spring setting that lies in the Parvati Valley in the Northern hemisphere of the great Himalayas (31°59’34" N, 77°30’35" E, altitude 2978) at district Kullu, Himachal Pradesh, India (Figure 1a-1b). The Parvati valley emerges from the confluence of rivers Parvati and Beas adjoined the hot water springs in the valley, including the hottest at Manikaran (Sangwan et al., 2015) have long been associated with high sulfur contents characterized by the distinct smell from sulfur dioxide or hydrogen sulfide gases escaping into the air (Shirkot and Verma 2015). Hot water is continually discharged from a major outlet at Khirganga from where it flows in a stream and deposits sulfur upon white microbial mats over the sediments along its course (Sharma et al., 2004). For this study, samples from all three habitats *viz.* microbial mats, sediments and water were collected proximal to the major opening (KgM1, KgS1, KgW1) and from a distance of 10m (KgM2, KgS2, KgW2) shown in Figure 1c.

**Figure 1:**
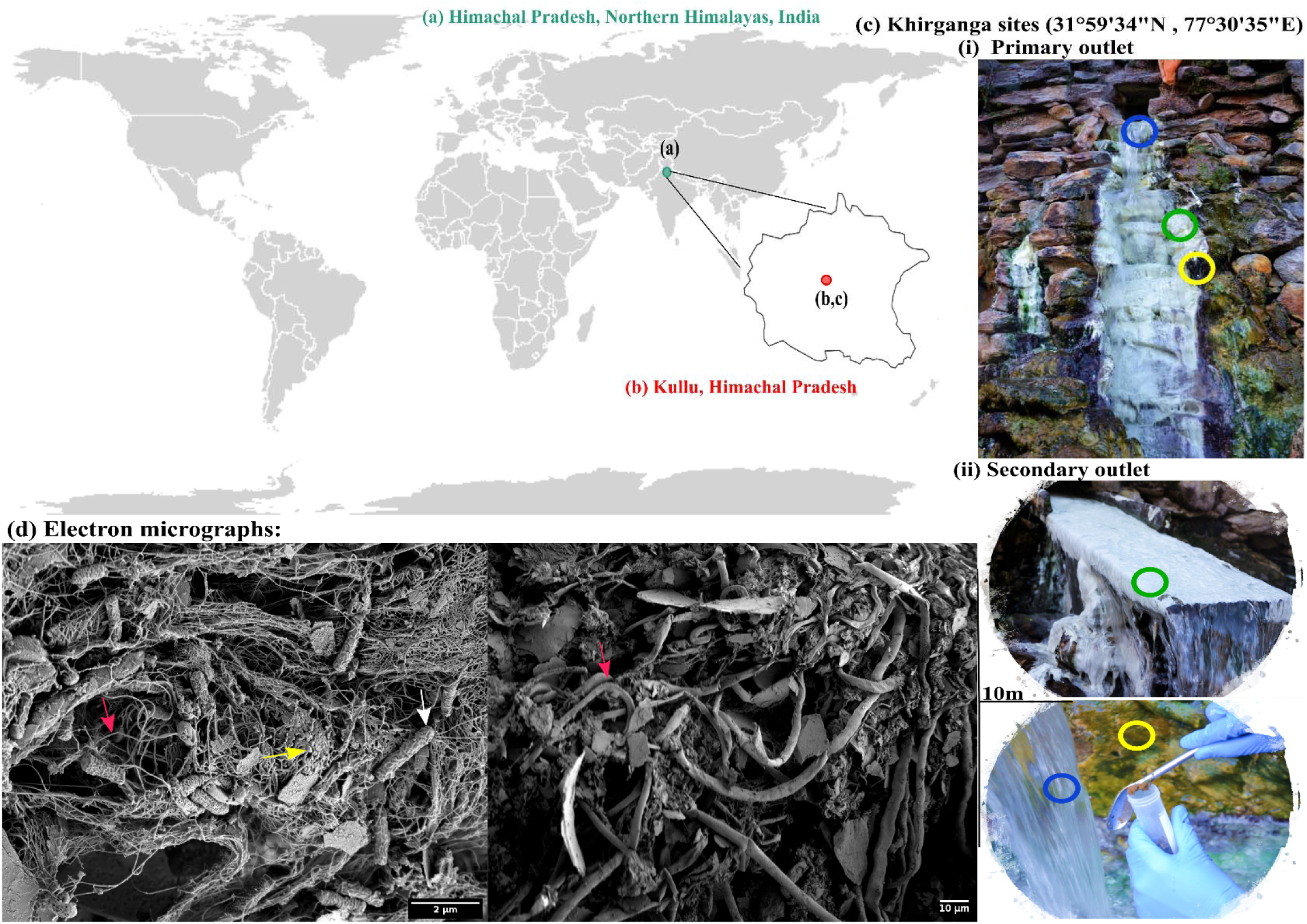
a) Geographical location of Himachal Pradesh, Northern Himalayas, b) Khirganga hot spring as shown in the maps are located in Parvati valley in Kullu district of Himachal Pradesh, India, c) the different habitats from where the samples were collected are shown: microbial mat (green), sediment (yellow) and water (blue). Samples were collected in replicates from two outlets located 10m distance apart d) the scanning electron micrographs of different habitat samples shown with arrow demarcating the filamentous (cyanobacteria), cocci-shaped and rod-shaped bacteria (SRB) in pink, yellow and white respectively.

Using helium-ion microscopy, we dissected the microstructure of the niches and were able to visualize cellular structures on complex sample matrices. The microbial diversity was visualized as numerous filamentous structures in microbial mats and sediments that resembled Cyanobacteria. Cyanobacteria are widely distributed in mats and sedimentary deposits of thermal springs (Gemerden et al., 1993; Podar et al., 2020). In addition, rod and cocci shaped cells of varying sizes were also observed in the sample matrices that providing a visual insight into the microbial diversity at this mesothermic site (Figure 1d).

### Physicochemical and Elemental Analysis

The *in-situ* measures of water temperature were from 59 at the outlet to 55 at 10m distance (Table 1). Sediments and microbial mat deposits had much lower temperature (42-45 ) than water. The pH of the hot spring water was 6.7 while sediments and mats were slightly acidic with pH 6.1 and 6.3 respectively. Thus, all three habitats were recorded to be mesothermic. The physicochemical composition of the hot spring is dominated by anions of chloride (up to 11024 mgL^-1^) and sulfate (up to 10079 mgL^-1^) while ions of calcium and potassium were abundant (Table 1; Supplementary File: 1). Importantly, sulfates (SO_4_^2-^) concentration in microbial mats and sediments were higher (9529 ± 313.29 mgL^-1^; 10079 ± 863.29 mgL^-1^, respectively) and exceeded the limit of 8000 mgL^-1^ standardized by Environment Protection Act (EPA, 2001) and also found to be exceeded the limit of 53 mgL^-1^ in surface waters (79.82 ± 1.85 mgL^-1^) (EPA, 2001; Table 1). The chlorides (1456.77 ± 367.27 mgL^-1^), manganese, sodium and silicon constituents in the hot spring waters were surpassing the normal average concentrations of 250, 0.05, 200 and 4 mgL^-1^ respectively (EPA, 2001) in surface water samples (Supplementary File: 1). Among others, the predominant elements and minerals in water samples were aluminium, magnesium, copper, zinc and arsenic. The data signified that the hot spring waters have high concentrations of sulfates and chlorides that are characteristic of majority of other hot springs in the Himalayan ranges (Cinti et al., 2009).

**Table 1:** Physicochemical and elemental analysis of the niche samples. Values for microbial mats and sediments samples are given in (ug/g) and those of water samples are shown in ppm.

| Value for indicated sampling sites |  |  |  |
| --- | --- | --- | --- |
| Environmental data | Mat | Sediment | Water |
| Temperature (°C) | 42.4 ± 0.56 | 44.55 ± 1.06 | 57 ± 2.2 |
| pH | 6.2 ± 0.14 | 6.5 ± 0.14 | 6.7 ± 0 |
| Major ions/metals | (ug.g<sup>-1</sup>) | (ug.g<sup>-1</sup>) | (mg.L<sup>-1</sup>) |
| Cl<sup>-</sup> | 11024.88 ± 983.17 | 15547.08 ± 204.96 | 1456.77 ± 367.27 |
| SO<sub>4</sub><sup>2-</sup> | 9529.06 ± 313.29 | 10079 ± 863.29 | 79.82 ± 1.85 |
| Na<sup>+</sup> | 976.88 ± 64.83 | 991.47 ± 7.88 | 1338.52 ± 167.97 |
| Ca<sup>2+</sup> | 7529.71 ± 1034.68 | 5111.4 ± 588.21 | 30.27 ± 7.32 |
| K<sup>+</sup> | 3681.66 ± 886.65 | 7743.91 ± 412.12 | 41.56 ± 9.03 |
| Al | 7205.41 ± 1685.37 | 22011.1 ± 2312.9 | 0.05 ± 0 |
| Mg | 3528.62 ± 735.04 | 8204.28 ± 964.02 | 8.83 ± 0.02 |
| Si | 3.22 ± 0.66 | 5.28 ± 0.42 | 31.5 ± 1.36 |
| Mn | 1149.7 ± 178.85 | 1219.29 ± 13.93 | 0.34 ± 0.03 |
| Cu | 14.4 ± 2.62 | 27.23 ± 1.71 | <0.01 |
| Zn | 27.44 ± 6.36 | 67.35 ± 0.47 | <0.01 |
| As | 3.9 ± 0.64 | 6.87 ± 0.22 | <0.01 |
| Ag | 5.51 ± 3.7 | 5.1 ± 0.03 | <0.01 |
| Pb | 17.74 ± 2.54 | 11.4 ± 1.27 | <0.00 |

In general, hydrogen sulfide account for the sulfur present in the underground geothermal waters originating from pyrites or leaching of other sulfides by deep hypothermal waters (Picard *et al.,* 2016). Sulfide (S^2-^) is oxidized to sulfate (SO_4_ ) as the water rises to the surface and under mild oxidizing conditions, sulfide is only oxidized to sulfur or sulfur dioxide (Picard *et al.,* 2016; Wu *et al.,* 2021). The results provided pertinent information on the geochemical composition of the three habitats to be correlated with the microbial diversity and community functions. More importantly, high concentrations of sulfate ions in microbial mats and sedimentary deposits supported the hypothesis of a key role of the bacterial sulfur cycle in sustaining the microbial community at the hot water spring.

### Metagenomic DNA Sequencing and Assembly

A large metagenomic dataset was obtained from sequencing having sufficient no. of reads sized up to ∼18Gb for each sample. We retrieved a total number of reads ranging between 110,861,650 to 152,895,302 in all samples which were assembled into 180,849 - 519,194 (>200 bp) contigs. After assembly, the metagenomes sizes varied between 329 and 600 Mbp. A summary of characteristics of the datasets and assembled metagenomes is provided in Table 2. The alpha diversity estimated as the Shannon diversity indices ranged between 2.5 and 3 (Table 2; Supplementary File: 2). The alpha diversity was higher in the water samples than microbial mat and sediments as has been reported previously (Ghilamicael *et al.,* 2017). The species richness as rarefaction curves obtained for all samples attained a plateau indicating optimum sequencing and sampling of a reasonable number of species for all metagenomes. The Bray-Curtis index calculated and plotted using NMDS demonstrated a significant difference in the beta-diversity of all three habitats at phyla level (PERMANOVA; P < 0.01). Abundance of class γ*-Proteobacteria* in microbial mats significantly distinguished latter from the other two habitats. Similar results have been reported in previous studies (Selvarajan et al., 2018, Pohlner et al., 2019). Communities in sediment and water samples varied from each other majorly due to differences in the abundance of *δ-Proteobacteria*.

**Figure 2:**
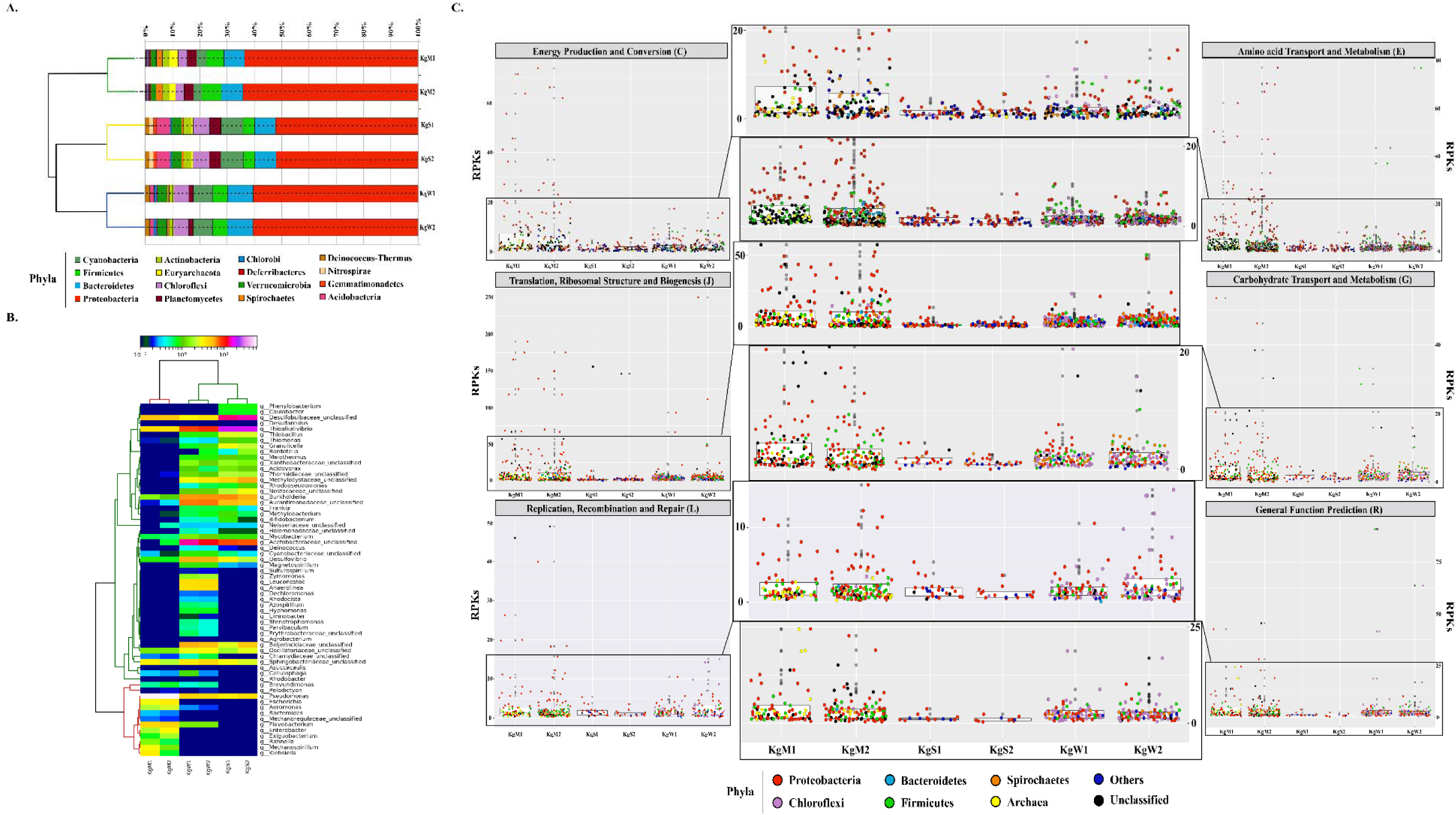
Relative abundance of phylum and genera. A) The stacked bar representation shows the dominating phylum in all three habitats. The dendrograms show hierarchical clustering between species and samples. B) The abundant common genera *Pseudomonas* (20–60%), *Desulfobulbaceae_unclasssifed* (15– 20%), *Burkholderia* (10-20%), *Desulfovibrio* (5-10%) and *Thioalkalivibrio* (5–7% are shown here relative abundance x log_x_ scale. C) Taxa based functional profiles demonstrating the major phylum contributing towards COG subsystems and percentage of proteins annotated within each COG category for the habitat sites. The distribution of the RPKs were mapped in accordance to the member abundance in the habitats. Dots were colored according to the phylum.

**Table 2:**
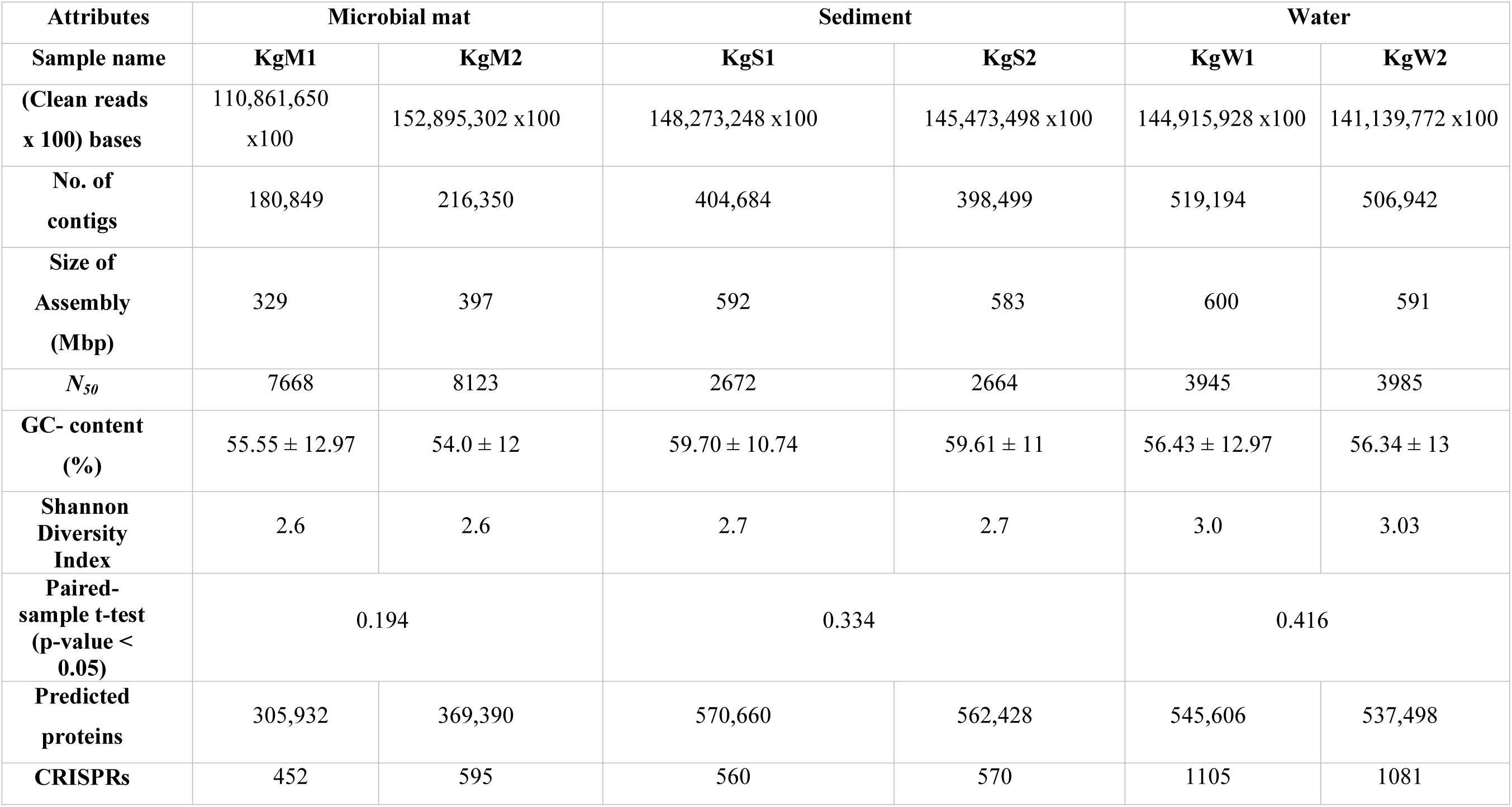
Characteristics of sequenced datasets generated and assembled metagenomes obtained for each sample from different habitats.

| Attributes | Microbial mat |  | Sediment |  | Water |  |
| --- | --- | --- | --- | --- | --- | --- |
| Sample name | KgM1 | KgM2 | KgS1 | KgS2 | KgW1 | KgW2 |
| (Clean reads x 100) bases | 110,861,650 x100 | 152,895,302 x100 | 148,273,248 x100 | 145,473,498 x100 | 144,915,928 x100 | 141,139,772 x100 |
| No. of contigs | 180,849 | 216,350 | 404,684 | 398,499 | 519,194 | 506,942 |
| Size of Assembly (Mbp) | 329 | 397 | 592 | 583 | 600 | 591 |
| $N_{50}$ | 7668 | 8123 | 2672 | 2664 | 3945 | 3985 |
| GC- content (%) | $55.55 \pm 12.97$ | $54.0 \pm 12$ | $59.70 \pm 10.74$ | $59.61 \pm 11$ | $56.43 \pm 12.97$ | $56.34 \pm 13$ |
| Shannon Diversity Index | 2.6 | 2.6 | 2.7 | 2.7 | 3.0 | 3.03 |
| Paired-sample t-test (p-value < 0.05) | 0.194 |  | 0.334 |  | 0.416 |  |
| Predicted proteins | 305,932 | 369,390 | 570,660 | 562,428 | 545,606 | 537,498 |
| CRISPRs | 452 | 595 | 560 | 570 | 1105 | 1081 |

### Dominance of Sulfur oxidizing bacteria (SOB) and sulfate reducing bacteria (SRB) in microbial consortia

Bacteria belonging to 15 different phyla dominated the microbial communities. The average percentage relative abundances of major phyla in the three habitats shown in parentheses in the order microbial mat, sediment and water as follows: *Proteobacteria* (62.1%, 50.5%, 58.7%), *Bacteroidetes*, *Firmicutes*, *Cyanobacteria, Planctomycetes*, *Chloroflexi* (Figure 2A). Species belonging to phylum *Proteobacteria* are found in varied temperature ranges which results in their dominance in various hot springs (De León et al., 2013; Singh et al., 2016; Saxena et al., 2017) and disproportionation of sulfur compounds is mainly carried out by SRMs of Proteobacteria (Finster, 2011), Besides, *Actinobacteria*, *Spirochaetes*, *Verrucomicrobia, Acidobacteria, Deinococcus-Thermus, Deferribacteres, Chlorobi, Gemmatimonadetes* and *Nitrospirae* were also detected with relative abundances less than 3%. Archaea superkingdom was represented by methane-producing phyla *Euryarchaeota* (2.5% in mat, 0.6% in sediment, 0.4% in water) and *Thaumarchaeota* which accounted for 0.5% of all classified sequences. These archaeal members are well known for actively carrying out ammonia-oxidation, nitrification and carbon fixation in marine and terrestrial environments in hot spring environments (Eme *et al.,* 2013; de la Torre *et al.,* 2008; Hatzenpichler *et al.,* 2008). Among the three habitats, microbial diversity profiles of water and sediments were more similar compared to those of microbial mats.

Highest genus level diversity was revealed in sediment (n=196) followed by water (n= 132) and mat (n= 63). Top 50 genera in all habitats were plotted (Figure 2B). Microbial mats were dominated by *Pseudomonas* (61.8%) followed by unclassified genera of family *Desulfobulbaceae* (SRB; 5.2%) and *Flavobacterium* (4.3%). Among the abundant genera in the microbial mats were *Aeromonas, Flavobacterium, Exiguobacterium, Enterobacter, Escherichia* and *Klebsiella* (Figure 2B). These taxa are often found associated with mat deposits and are active biofilm producers that are thought to use adherence to the surface as a strategy to survive, evolve and to cope with various abiotic stresses at such extreme habitats (López et al., 2006; Wright et al., 2013). On the other hand, sediment habitats were found to be enriched in *Thioalkalivibrio* (SOB; 18.9%), *Desulfobulbaceae* (SRB; 14%), *Burkholderia* (6.3%) and unclassifed genera of families *Acetobacteraceae* (8%). Hot spring waters with highest diversity of bacterial genera were dominated by *Thioalkalivibrio* (9%), *Burkholderia* (6.5%), *Desulfovibrio* (SRB; 5.9%). Other genera with less than 6% abundances in all three habitats were also detected as shown in Figure 2B.

One important observation was the dominance of sulfur-oxidizing bacteria (SOB) and sulfate-reducing bacteria (SRB) in all three habitats (Agostino and Rosenbaum, 2018; Nagar et al., 2021). SOB and SRB are usually categorized as lithoautotrophs that play key microbial role in biogeochemical cycling of sulfur in various habitats. SOB oxidize the reduced sulfur compounds such as hydrogen sulfide (H_2_S), elemental sulfur (S^0^), sulfite (SO_3_^-2^ ), thiosulfate (S_2_O_3_^2-^), and various polythionates (S_n_O_6_^2-^ or -S_n_O^6-^ ) into sulfate (SO_4_^-2^ ). On the contrary, SO_4_ can serve as an electron acceptor of SRB under anaerobic conditions, and they reduce the SO_4_^-2^ and other oxidized sulfur compounds (S_2_O_3_^2-^ , SO_3_^-2^ , S^0^) into H_2_S (Agostino and Rosenbaum, 2018). The abundance of SOB such as *Thioalkalivibrio* and *Burkholderia* as well as SRB such as *Desulfobulbaceae unclassified* and *Desulfovibrio* in spring is not surprising as high levels of sulfate dominate the site and relative abundance of these bacteria provide evidence of an active S cycling mediated by microbial communities. We identified the sequences producing significant alignments from the nr database for taxonomic confirmation and assigned each sequence that could be classified up to genus level to either SOB or SRB (Figure 4A, Supplementary File: 5). The taxonomy and phylogenetic topologies are discussed in detail in the next section.

**Figure 4:**
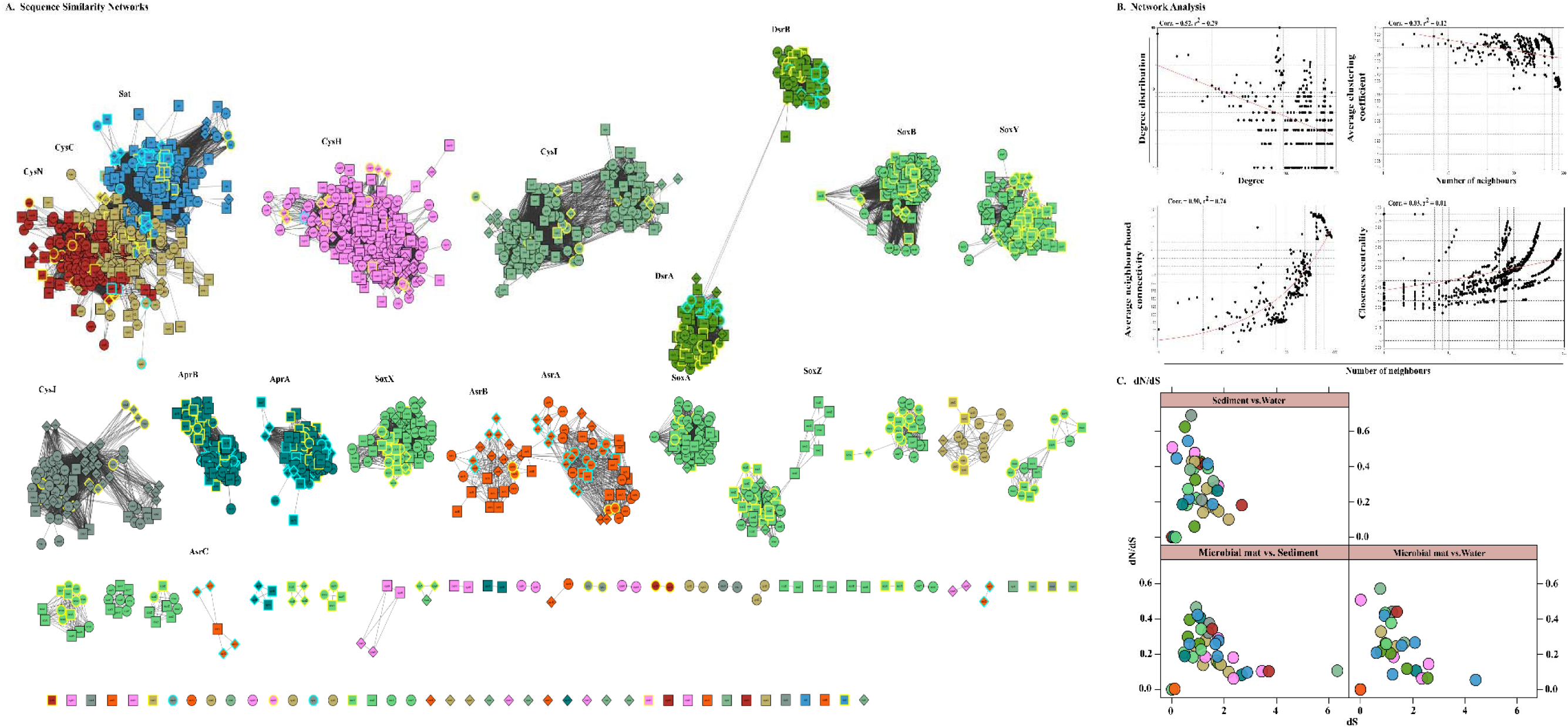
Sequence similarity network analyses. A) Diversity of sulfate reduction genes of both assimilatory and dissimilatory pathways in microbial mat (diamod-shaped), sediment (square) and water (spherical) habitats visualized in cytoscape v3.7.1. Highlighted only the classified taxa, where color cyan belong to SRBs and yellow to SOBs. The network was set at threshold e-value cutoff of 1e-30 and nodes represent sequences connected through edges if the similarity exceeds the cutoff score. Here, the clusters and isolated nodes were showing the conserved pattern and diversified pattern of the proteins significantly playing an important role in sulfate reduction, B) Topological properties of the similarity networks: degree distribution, average clustering coefficient, average neighborhood connectivity and closeness centrality are plotted against the number of neighbors. The power law fit curves are shown within each graph. C) Habitat vs. habitat dNdS values of all S cycle genes were estimated and plotted using xyplot in R (R Development Core Team, 2011).

### Metabolic functions of the community and sulfur metabolism among the three sampling sites

The major gene families inferred to be abundant across all three habitats were assigned to *Proteobacteria* followed by *Chloroflexi*, *Firmicutes*, *Bacteroidetes* and *Spirochaetes* as reflected from the RPKs (reads per kilobase) in the metagenomes. These gene families were then regrouped as clusters of orthologous groups (COGs) and the top functions were determined to be translation, ribosomal structure and biogenesis (COG: J), amino acid transport and metabolism (E), general function prediction (R), energy production and conversion (C), replication, recombination and repair (L) and carbohydrate transport and metabolism (G). These functions in microbial mats were carried out by *Proteobacteria* (COGs: J, E, C, G) and unclassified bacteria (COGs: R, L); in sediments by unclassified bacteria (COG: J) and *Proteobacteria* (COGs: E, R, C, L, G) and in water by *Proteobacteria* (COGs: J, C), *Firmicutes* and *Chloroflexi* (COGs: E, R, G, L) (Figure 2C; Supplementary File: 3).

The ORFs that were categorized on the basis of KEGG categories were mapped onto the metabolic functions and the pathways that could be reconstructed with more than 60% completeness were used to define the metabolic potential of the habitats. Based on this criterion, we studied the top 50 functional pathways of each habitat and identified the core functions (n=37 pathways) of the communities that included the common pathways for metabolism of nucleotides, carbohydrates and amino acids (Figure 3A, Supplementary File: 4). In addition, we determined differentially abundant pathways in each habitat: microbial mats (n=7), sediments (n=7) and water (n=2). Sediments were enriched in functions such as glycosaminoglycan degradation, metabolism of cyanoamino acid and riboflavin as well as degradation of nitrotoluene, geraniol, lysine and fatty acids while communities in water showed abundance of fatty acid and novobiocin biosynthetic pathways (Figure 3A). The community functional profiles of sediment and water were more similar compared to those of microbial mats which may be due to the stratified layered organization of the mats which are different in sediment and water. The microbial communities in mats were optimized for metabolism of methane specifically, members of genus *Methanospirillum*, which were abundant in mats (Figure 2B). Methanogens are shown to compete more effectively with SRB in attached-growth systems such as those of mats in which kinetic and thermodynamic considerations are the dominant mechanisms regulating the competition (Raskin et al., 1996). Other metabolic pathways such as lysine biosynthesis, C5-branched dibasic acid metabolism, thiamine metabolism, pyrimidine metabolism, vitamin B6 metabolism and other glycan degradation were also found to be abundant in microbial mat.

**Figure 3:**
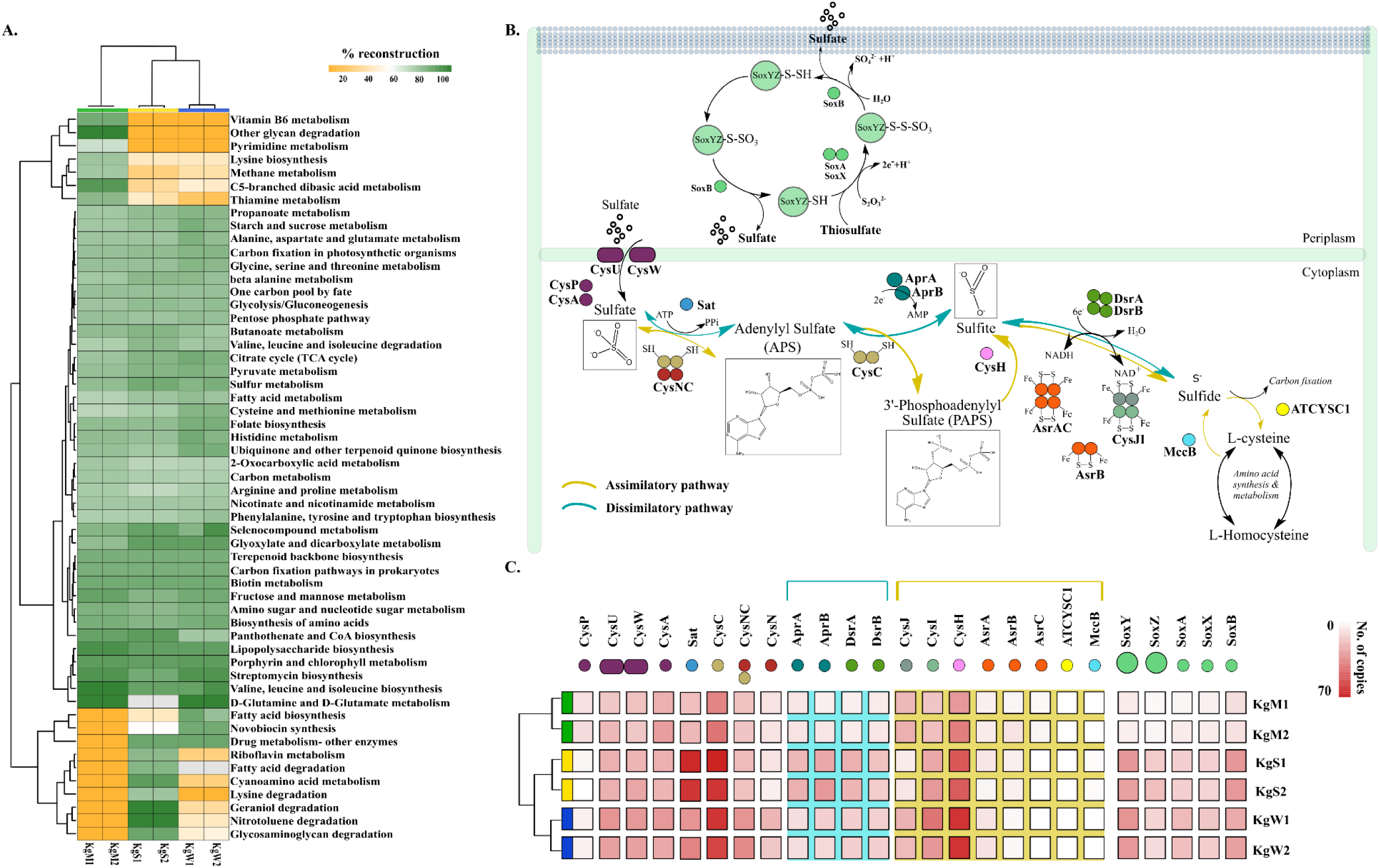
A) Reconstruction of top 50 pathways annotated using KEGG Automatic Annotation Server. Heatmap matrix representation and clustering was performed by using ‘pheatmap’ package (Kolde and Kolde, 2015) in R (R Development Core Team, 2011 http://www.R-project.org/). B) The sulfate reduction pathway involved a group of reductases, kinases and transferases with the product chemical structures generated through chemDraw7 and Inkscape v0.9 (Inkscape project, 2020). C) The gene copy number of both sulfate reduction and sulfide oxidation pathway that were partitioned in different habitats showed here using ggplot2 in R (R Development Core Team, 2011 http://www.R-project.org/).

The enriched diversity for sulfur oxidation and sulfate reduction as well as the geochemical analysis of sulfide rich habitats compelled us to mine the regulatory genes involved in the different pathways of S cycle. In natural system, the sulfur intermediates viz., sulfate (SO_4_^−2^), thiosulfate (S_2_O_3_^2−^), sulfite (SO_3_^−2^) or elemental sulfur (S^0^) are reduced by different bacteria through two different reduction processes, *viz.,* dissimilatory and assimilatory reactions (Vermeij and Kertesz 1999; Zavarzin 2008; Figure 3B). In dissimilatory reduction, SRB utilize three enzymes (ATP sulfurylase, APS reductase, and sulfite reductase) to reduce sulfate and produce toxic hydrogen sulfide (Agostino and Rosenbaum, 2018; Kushkevych et al., 2020). On the contrary, sulfate is assimilated into organic compounds under assimilatory process (Figure 3B; Kushkevych et al., 2020).

The S metabolism pathway could be reconstructed within a range of 76.31 - 84.21% which was maximum in sediment and minimum in microbial mat. Total 75 genes were responsible for sulfur metabolism present in all the samples with a mean copy number of 1580 ± 249.2. In order to gain insights into the sulfur oxidation as well as assimilatory and dissimilatory sulfate reduction potential across all the habitats, we mapped the TIGRFAM and Pfam (Supplementary File: 4) ids of the 25 associated genes (mean copy number of 980 ± 222.1) on to the ORFs and copy numbers of these S cycling genes involved in sulfur oxidation, sulfide oxidation and sulfate reduction (as described in Figure 3B) were estimated. Thiosulfate ions are fused to the carrier complex of sulfur-oxidizing proteins *soxYZ* (*soxY*=145, *soxZ*=71), while L-cysteine S-thiosulfotransferase (*soxX*=76, *soxA*=86) and S-sulfosulfanyl-L-cysteine sulfohydrolase (*soxB*=145) mediate the hydrolytic release of reduced sulfur ions from sulfur-bound *soxYZ*. Besides, the dissimilatory sulfite reductase encoding genes: *dsrA*=95, *dsrB*=83 and anaerobic sulfite reductase subunits *asrA*=44, *asrB*=30, *asrC*=9 reduces sulfite to sulfide (Supplementary File: 4). The other 15 genes for reduction of sulfate ions included solute binding protein (*cysP*=37), ATP binding protein (*cysA*=168), transport system permease proteins (*cysU*=158, *cysW*=154), ATP sulfurylase (*sat*=254), transferases (*cysC*=348, bifunctional *cysNC*=155, *cysN*=75), phosphoadenosine phosphosulfate reductase (*cysH*=314), adenylylsulfate reductase subunit A and B (*aprA*=88, *aprB*=104), sulfite reductase flavoprotein alpha-component (*cysJ*=111), sulfite reductase hemoprotein beta-component (*cysI*=180), homocysteine desulfhydrase (*mccB*=6), and cysteine synthase (*ATCYSC1*=4). Figure 3C shows the comparative abundances of these proteins in the three habitats. Here, the results revealed that the microbial reduction of sulfate occurs largely through the dissimilatory pathway carried out by dissimilatory sulfite reductase (DsrAB) as compared to assimilatory sulfite reductase (AsrABC) mediated reduction. In nature, the dissimilatory pathway is shorter and thus, preferred route of microbial sulfate reduction (Kushkevych et al., 2020).

### Diversification, interactions and evolution of key enzymes involved in sulfur oxidation and reduction

To study the diversity and evolution of the key enzymes of SOB and SRB communities in this environmental biosystem, we employed a two-step strategy of comparing similarities of all sequences in a pairwise fashion through sequence similarity networks (SSN) analysis and further estimated the measures of the rates of non-synonymous to synonymous substitutions in their orthologous proteins between each pair of habitats. SSN effectively resolves the pairwise similarities of each sequence (node) with every other sequence of an enzyme or a group of enzymes for a pathway such that any two nodes are connected by edges only if they share sequence homology above a certain cutoff (here e-value of 1e-30). Thus, SSN provides for an accurate placement of a sequence among its putative homologs (Talwar et al., 2020).

Here, we examined the diversity among 19 key S substrates oxidizing and reducing proteins determined from the communities as discussed in Figure 3C, except membrane permeases (CysPUWA) and genes for cysteine synthesis (MccB, ATCYSC1). In total, we retrieved 2413 protein sequences (mean; M=254, S=480, W=472) denoted by nodes in SSN. The network was organized into 88 connected components including 46 isolated nodes with an average clustering coefficient of 0.84. The connected components were represented by the homologous and heterologous clusters depending on whether they were constituted by the same gene or a number of different genes involved in a pathway, respectively. The number of connected components formed through SSN analysis of S metabolic proteins distributed into gene clusters and the isolated nodes denoted the diverging and highly diverged sequences, respectively (Figure 4A). Hence, we looked into these components to study the diversity of each gene that were distributed as shown in Table 3. The proteins CysNC, CysH, CysI and CysJ catalyse important steps and act as co-factors for the assimilatory sulfite reductase (AsrABC) which were all found to be diverging with many isolated nodes and loosely connected components. The enzymes for S oxidation (Sox) were also found to be diverging as observed from loosely formed clusters. On the other hand, all sequences of the key enzyme of dissimilatory pathway, DsrAB, formed only one connected component which suggested that they might be under convergent evolution at this site (Figure 4A). Further, we compared the distribution of diverged sequences that could be separated as isolated nodes and found that hot spring sediments harbored a high diversity of these enzymes (n=23) in comparison with microbial mats (n=12) and water (n=11).

**Table 3:**
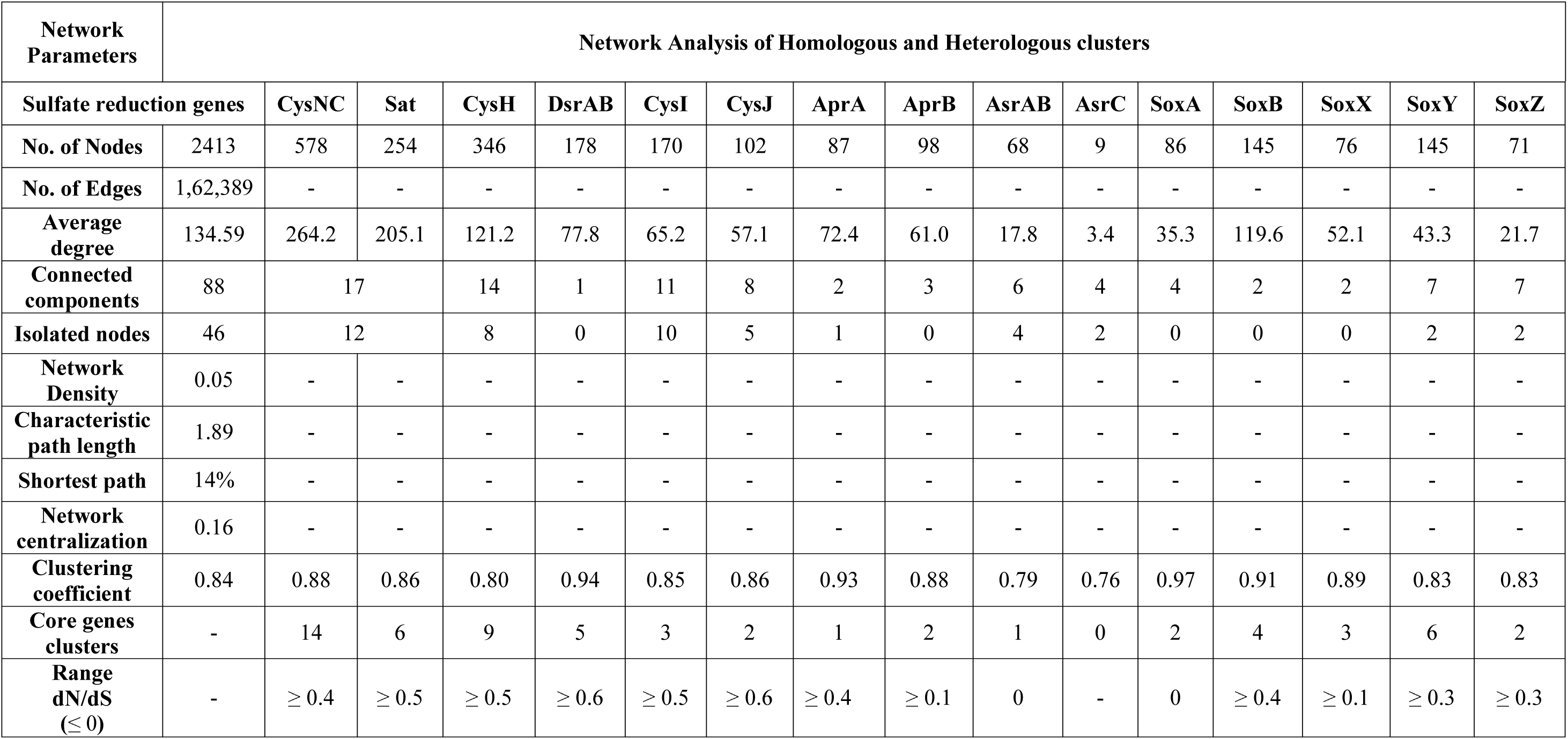
Attributes of the sequence similarity networks and dN/dS analysis of S metabolism genes.

Based on sequence similarity, majority of the AsrABC genes that were taxonomically related to SRB could be distinguished from the rest of the sequences that were not identified as either SOB or SRB. Thus, assimilatory reduction was diverged among the SRB communities while those for dissimilatory pathway were rather conserved at the site. Previously, sequence comparisons have confirmed that dissimilatory sulfite reductase, Dsr, to be highly conserved enzyme that could serve as marker gene for SRBs (Loy et al., 2009). The DsrAB and AprAB enzymes were contributed by both SOB and SRB with syntrophic interaction which suggests for the presence of reverse dsr (rdsr) mediated oxidation of S substrates in addition to dissimilatory reduction. Here, the inherent complexity of sulfur-based metabolic network revealed that there are controlled mutation rates in dsrAB genes in presence or increased selective pressure of contamination and extreme conditions. The syntrophy of SOB and SRB prevailing in anoxic and anaerobic conditions governs the dissimilatory sulfur metabolism (oxidation and reduction parallelly) and indirectly promoting the growth of diverse microbes in this natural ecosystem (Bhatnagar *et al.,* 2020).

Through network analysis, we observed that, the node degree distribution estimated to be decreasing with increasing protein quantity (correlation=0.52, r^2^=0.29), average neighborhood connectivity within the networks interpreted as function in *k* was increasing and positively correlated (correlation=0.90, r^2^=0.74) (Figure 4B). Furthermore, closeness centrality curve that measures closeness between nodes was unable to reach the bench top (correlation=0.03, r^2^=0.01), might be due to maximum number of connected components and less sequence homology. So, we also analyzed each protein cluster individually by using network analysis, 178/2413 nodes of the network DsrAB protein cluster found to be conserved showed higher clustering coefficient values (0.94), followed by AprA (0.93), AprB, CysNC (0.88) and others (Table 3). A relatively high diversity of the sulfate reduction proteins in all three segments of this habitat unveiled the high nutritional demands and efficiency of the microbes towards uptake of a wide range of structurally and chemically diverse amino acid side chains from environment (Talwar et al., 2020).

The evolutionary selection pressures on these genes were studied through estimation of dN/dS values calculated for a subset of conserved gene sequences in all three habitats (Figure 4C). The number of core genes and the range of dN/dS values identified for each gene are shown in Table 3. The CysJ and CysI genes were found to be under moderate selection pressures with dN/dS values in the range 0.4-0.7 (Figure 4C, Supplementary material: 6). The results supported the observation as these enzymes code for important co-factors for the assimilatory sulfite reductase (AsrABC) that were found to be diverging in through SSN analysis. Therefore, the microbial genes for assimilatory reduction pathway are diversifying under moderate selection pressures.

### Divergence, phylogeny and structural relationships of DsrAB and AsrAB

The enzymes catalyzing the reductive (dissimilatory) or oxidative (rDSR) transformation between sulfite and sulfide appear to be related with respect to their subunit composition and catalytic properties (Loy et al., 2009). The dsr genes have been characterized from bacterial as well as archaeal domains (Chang et al., 2001; Grim et al., 2011; Colman et al., 2020). However, their evolution in these domains has long remained a subject of discussion. Our preliminary results showed that both subunits of dsr genes (dsrA=79, dsrB=72) corresponds to about 70 newly identified organism for both oxidation and reduction processes (Supplementary File 8). Through RAxML phylogenetic analysis, it can be confirmed that the dsrAB genes have been introduced in most of the newly identified members by a multiple independent Lateral Gene Transfer (LGT) (Anantharaman *et al*., 2018). Importantly, organisms from *Acidobacteria*, *Candidate division Zixibacteria*, *Chloroflexi* and β*-Proteobacteria* form completely novel lineages other than known DsrAB clusters identified through RefSeq *nr* protein database with accession numbers (National Centre for Biotechnology Information; Supplementary File: 5). The result of the phylogeny suggested that an increase in substitution rate in both subunits of DSR may have occurred on the branch connecting δ*-Proteobacteria* to all other taxa as observed from the branch lengths. The different rates of substitution of the two DSR subunits has hitherto only been reported in δ*-Proteobacteria* lineage (Wagner et al., 1998). Hence, the dsr from sulfate reducers formed a separate cluster, with sequences from *Desulfarculus, Desulfocarbo, Desulfarcinum, Thermodesulfobacteria, Syntrophobacter, Desulfomonile, Desulfovibrio, Desulfatirhabdium* and *Desulfobacteriaceae* in both DsrA and DsrB phylogenies and additionally, *Dissulfuribacter* in DsrB (Supplementary File 8). We proposed that these organisms with newly identified lineages of dsr genes involved in sulfite/sulfate oxidation and reduction likely serves an important control on sulfur cycling on terrestrial subsurface. The divergence of dsrAB between unrelated taxa could be driven through combination of speciation, functional diversification and LGT. Also, there is equal possibility of non-functionality of the genes in these taxa (Loy *et al.,* 2008; Anantharaman *et al.,* 2014).

Through time-scale evolutionary phylogeny of the sequences of DsrAB and AsrAB identified from the metagenomes, we determine the most basal and earliest evolved lineages involved in dsr and rsdr pathways (Figure 5; Supplementary File: 7). Our results suggested that both the subunits of the oxidative type reverse-DSR evolved much earlier than the reductive type DSR subunits (Figure 5). The templates of these *dsrAB* genes have potential to study the genotypic and phenotypic traits in SRPs and the dissimilatory sulfur metabolism processes which will expand the gene-environment interaction mechanism. Also, prior analysis has proved that the evolution of *dsrAB* have been influenced by LGT only among major taxonomic lineages (Klein et al., 2001; Muller et al., 2015) but the findings here provide evidence of independent multiple LGT events distributed throughout the dissimilatory gene clusters. Currently, the time scale study of this site cannot produce evidence of the progenitor lineages, as the evolutionary history of dissimilatory reduction is complex and yet ascertain. Although it had provided information of the earliest lineage where sulfate/sulfite oxidation and reduction appeared.

**Figure 5:**
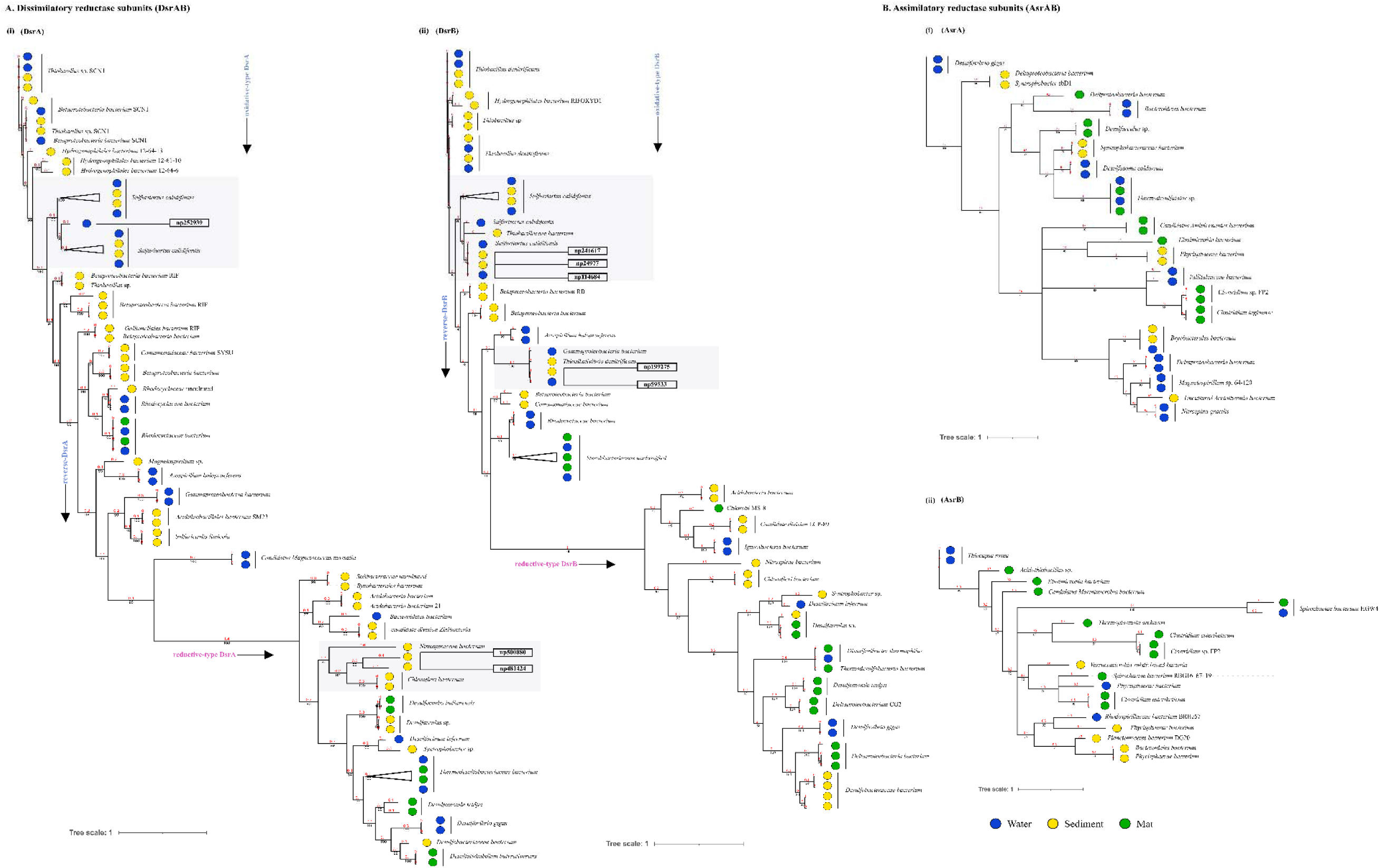
Divergence estimation over time: Reconstruction of the phylogenic tree of optimized full length dissimilatory sulfite reductase and assimilatory sulfite reductase subunits in three habitats using PhyloBayes with the CAT-GTR model; The highlighted squares consist of clades with proteins that were remained unclassified through *nr* database. A. (i) Among, 78 DsrA nodes that showed here the earliest evolution of the rDsr oxidative proteins occured in *Thiobacillus* sp (ii) 72 nodes of DsrB proteins with similar results; B. (i) 38 nodes of AsrA (ii) 21 nodes of AsrB were also compared as a control for branch length shown here.

We used the phylogenetic analysis to further assign taxa to the sequences that showed similarity with yet uncultured bacteria and predicted their structures to gain insights into the more common phylogenetic ancestor of the two DSR subunits. Interestingly, these sequences of the oxidative type Dsr (rDSR) formed monophyletic clade with a more recently identified genus, *Sulfuritortus* in both DsrA and DsrB phylogenies. The genus of sulfur oxidizing bacteria currently has only one recognized species and is closely related to the members of genus *Thiobacillus* (Kojima et al., 2017). Our analysis revealed similar tree topologies with these unassigned sequences forming clade with *Sulfuritortus, Thiobacillus* and *Hydrogenophilales bacterium* in both DsrA (n=1) and DsrB (n=3). Although the sequences were similar to *Thiobacillus* phylogenetically, prediction of their structures revealed DsrA to be highly similar to that of *Desulfovibrio gigas* (TM score= 0.75; Analog TM= 0.86; C-score=0.25, 1.6, 1.58; Supplementary File 9) and structures of DsrB proteins aligned most closely with *Archaeoglobus fulgidus* (TM score= 0.94; Analog TM= 0.97; C-score=1.64; Supplementary File 9, 10). While the other DsrB subunit (n=2) formed clades with *Thioalkalivibrio* (β*-Proteobacteria)* also showing similarity to *A. fulgidus* (TM score= 0.94; Analog TM= 0.98; C-score=1.64).

In the reductive type Dsr clades, two DsrA sequences from uncultured bacteria formed clades with *Nitrospiraceae* and Chloroflexi bacteria (Figure 5). However, a consistent observation was seen in the similarity of all these structures of DsrA proteins with *Desulfovibrio vulgaris* (TM score= 0.94; Analog TM= 0.96; C-score= 1.58). Thus, we predict that the genus derives its two subunits of Dsr from different ancestors. A plausible explanation for this is observed in the previous reports of high sequence homology between the Dsr of *A. fulgidus* and *Desulfovibrio vulgaris* that suggest a common origin of archaeal and bacterial DSRs or their horizontal gene transfer (Karkhoff-Schweizer et al., 1995). In addition to homology in their sequences, the evolutionary distance separating the enzymes from *A. fulgidus* and *Desulfovibrio vulgaris*. For DsrB subunits, the archaeal and bacterial sequences were not particularly distant; such that the branches with structural homology to *A. fulgidus* were approximately the same length as branches leading to bacteria such as *Thiobacillus, Thioalkalivibrio* and *β-Proteobacteria bacterium* (Larsen et al., 1999).

## Conclusion

To date, the microbial community and functional dynamics including genes of sulfur hot spring in Himalayas are poorly understood. This study was designed to investigate the syntrophic interactions between SOB and SRB in sulfur contaminated spring. Through analysis of microbial mat and sediment deposits along with water samples, we traced the microbial diversity and evolutionary dynamics of genes involved in S cycling within the Khirganga ecosystem. The mesothermic hot spring have been composed of a diverse group of microbes (Bacteria and Archaea) and genotypes (*dsrAB*) that could be screened out as novel thermozymes that cannot be underestimated. From the results, it could be concluded that the microbial community functions were distinguished in microbial mat from water and sediment. These thermophilic microbial inhabitants played a very crucial role in understanding the metal toxification, ion exchange and biogeochemical cycling of the elements that later became acquainted with physicochemical and elemental analysis of the habitat. Here, the genomic repertoire suggested the ongoing specific adaptations to cope up with extreme values of chloride, sulfate ions and manganese, silica contamination in this ecological setting. The sulfur metabolic pathways are completed where inorganic sulfur compounds being the main source for sulfate reducing bacteria releasing toxicity in the form of sulfides (S^-^). The sulfate reduction profiling in all three habitats reveals dissimilatory sulfate reduction process (*dsrAB*) is active than assimilatory sulfate reduction (*asrAB*). Later, the genes involved in sulfur reduction/oxidation were classified and belong mostly to *Proteobacteria* with maximum homologous proteins classified in anoxygenic SOBs. In all sulfur disproportionation proteins, the sulfite reductase DsrAB proteins showed conserved behavior with 0/1 isolated nodes that have been signified as phylogenetic markers for SRBs. The evolutionary phylogenetic analysis showed that the oxidative rDsr were the earliest then the reducing Dsr which may predict that the condition with more sulfides oxidized in more sulfates, directing sulfate reducing bacteria (SRBs) to perform dissimilatory reduction later. Phylogenetic clades of DsrAB proteins showed unanimous distribution of taxa except the δ *Proteobacteria* which could be the reason for occurring LGT to other phyla. On the basis of structural alignments, the lineages with unclassified clades have shown different analogy in both Dsr subunits where DsrB derived from Archaea and DsrA are *δ-Proteobacteria* in origin at this meso-thermic niche.

## Availability of Data

All sequence data presented in this article is submitted and is available in the public repository of NCBI under Bioproject number PRJNA673998, experiment Biosample ID and SRA number are shown in Supplementary File 1.

## Acknowledgements

The sequence data were produced by the Beijing Genome Institute, China (BGI Tech. Solutions Co .Ltd , Hongkong) in collaboration with the user community. The authors are thankful for the use of the scanning electron microscope at the Centre for Chemical Microscopy (ProVIS) scanning at UFZ Leipzig, which is supported by European Regional Development Funds (EFRE-Europe funds Saxony) and the Helmholtz Association and The Energy and Resources Institute (TERI).

## Supplementary material

**Supplementary File 1:**
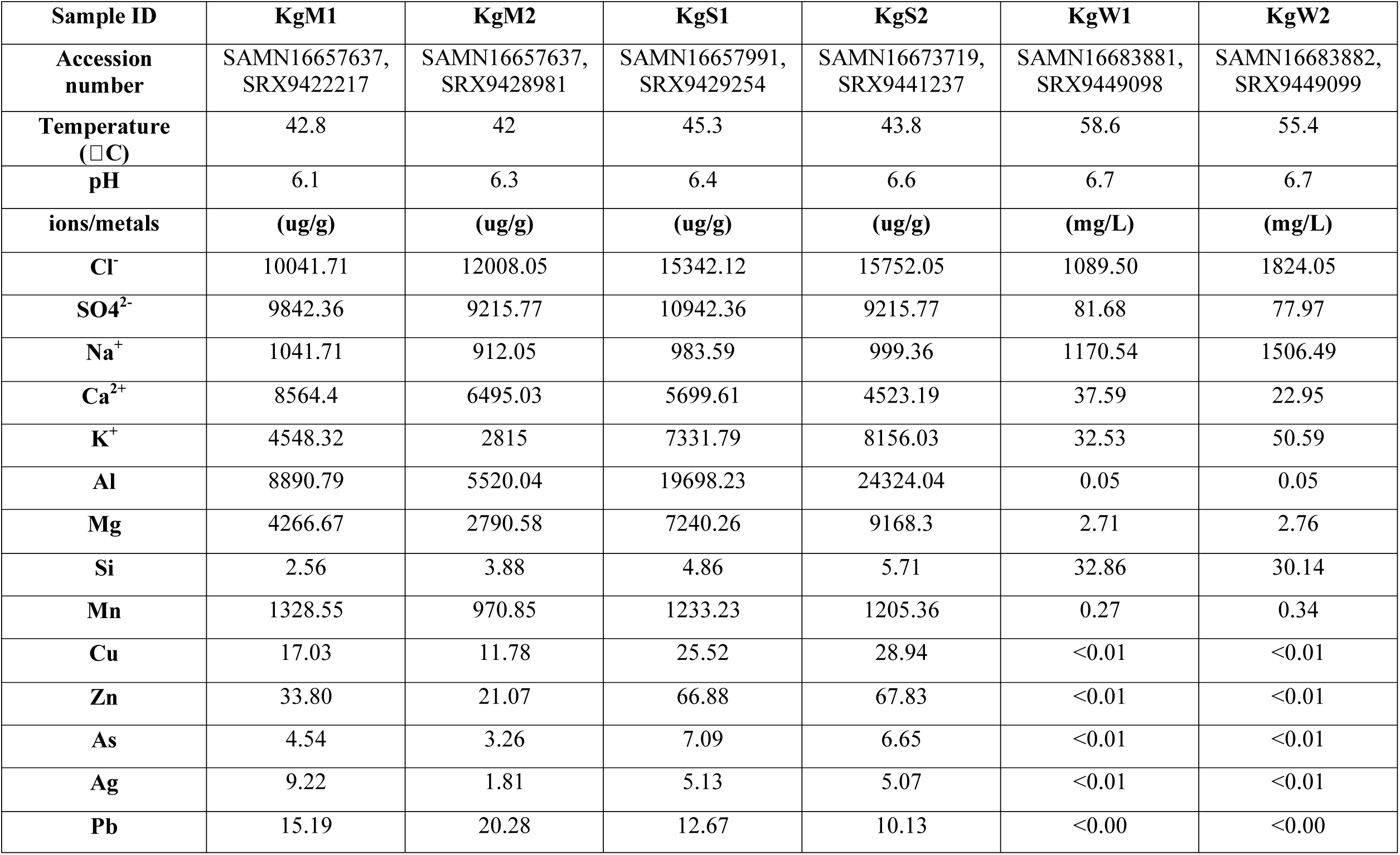
NCBI accession and in-situ measures of temperatures, pH and elemental analysis of each sample from three studied niches.

**Supplementary File 2:**
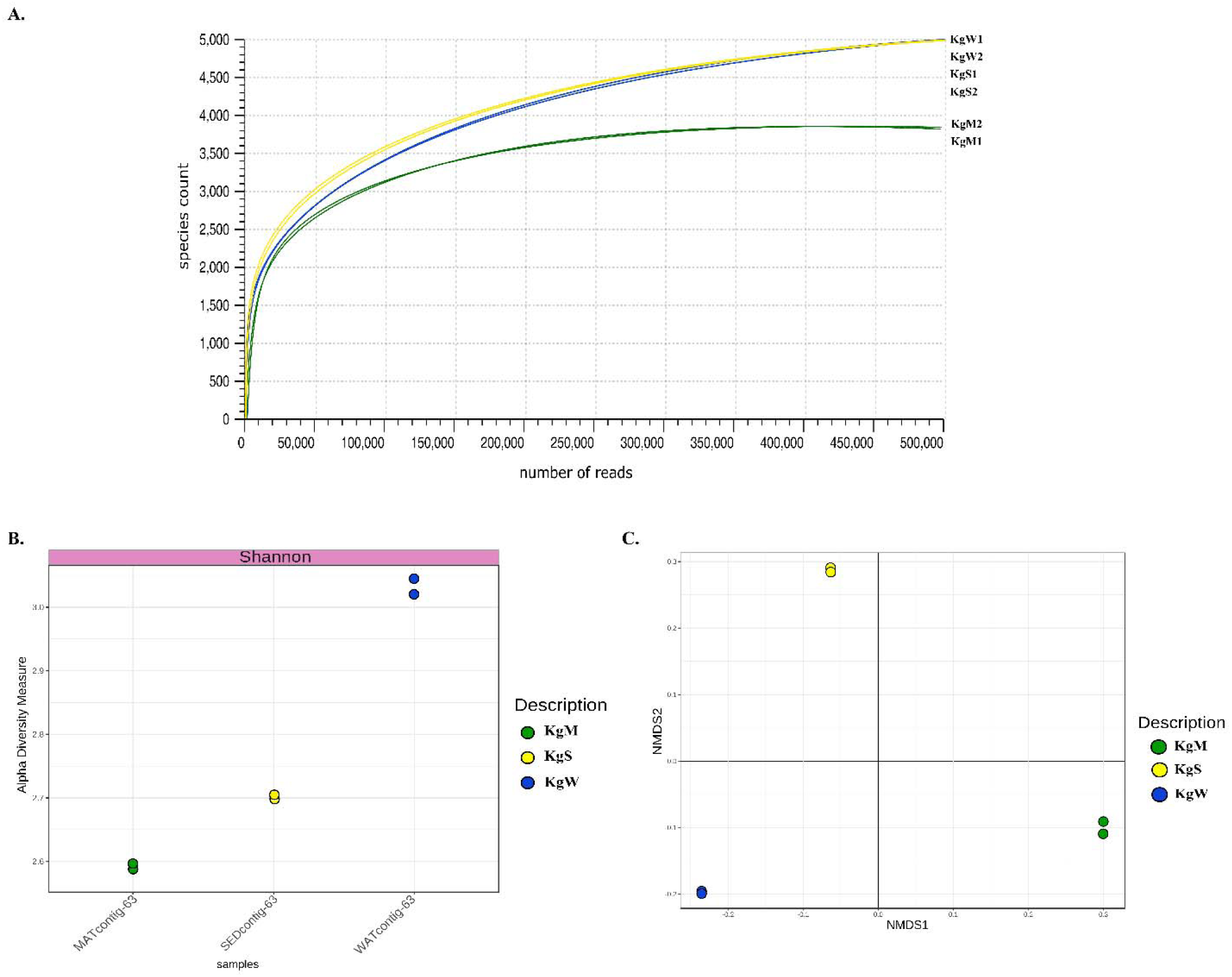
Rarefaction curves based on measures of alpha diversity species richness calculated for each of the six samples (B) The interquartile range of alpha diversity measures based on Shannon indices. (C) Non-metric multidimensional scaling (NMDS) plots based on Bray-Curtis indices of community dissimilarities in the six samples types (p <0.05, PERMANOVA).

**Supplementary File 3:** The list of all metabolic gene families in each sample (KgM1, KgM2, KgS1, KgS2, KgW1, KgW2) and the RPKs abundances of taxa (*Proteobacteria, Bacteroidetes* etc.) summarized in details.

**Supplementary File 4:** Detailed information of percentage reconstruction of metabolic pathways; Common and unique pathways in all six samples; Identification sources for 19 important proteins involved in sulfur disproportionation using TIGRfam and Pfam conserved domain database, ‘x’ indicates no hits found; Also, the copy number of 25 sulfur disproportionation proteins recorded.

**Supplementary File 5:** Details of taxonomic information of 2413 nodes of 19 sulfur cycle genes identified from nr NCBI microbial proteins database.

**Supplementary file 6:** Habitat vs. habitat dN/dS values of the sulfur cycling genes.

**Supplementary File 7:**
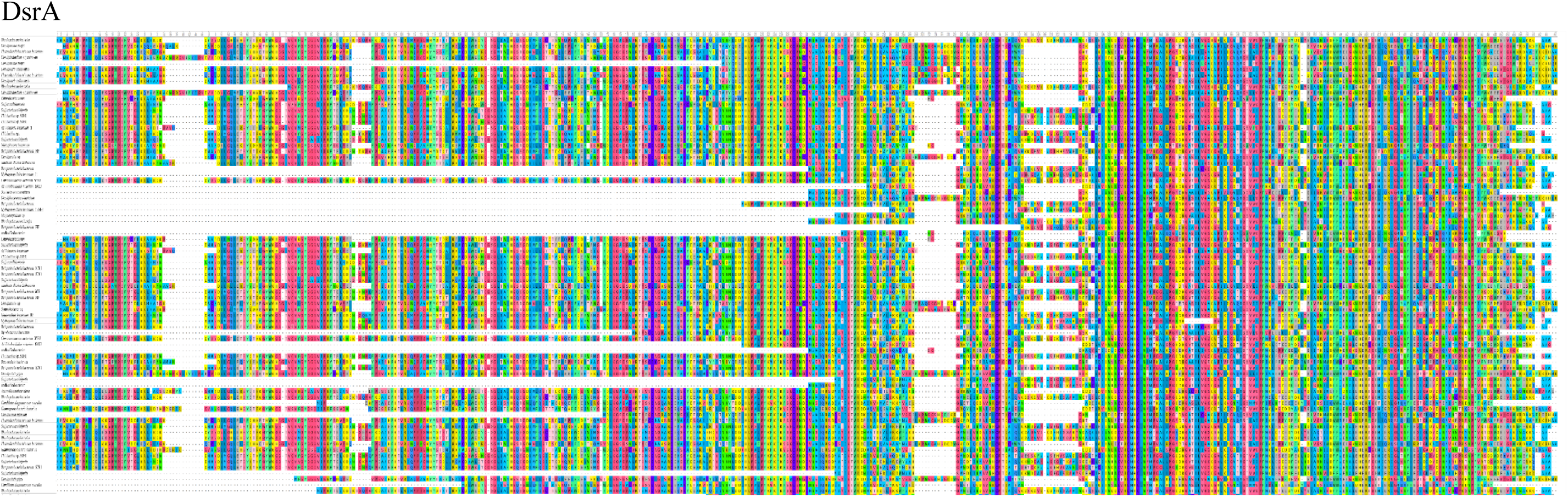
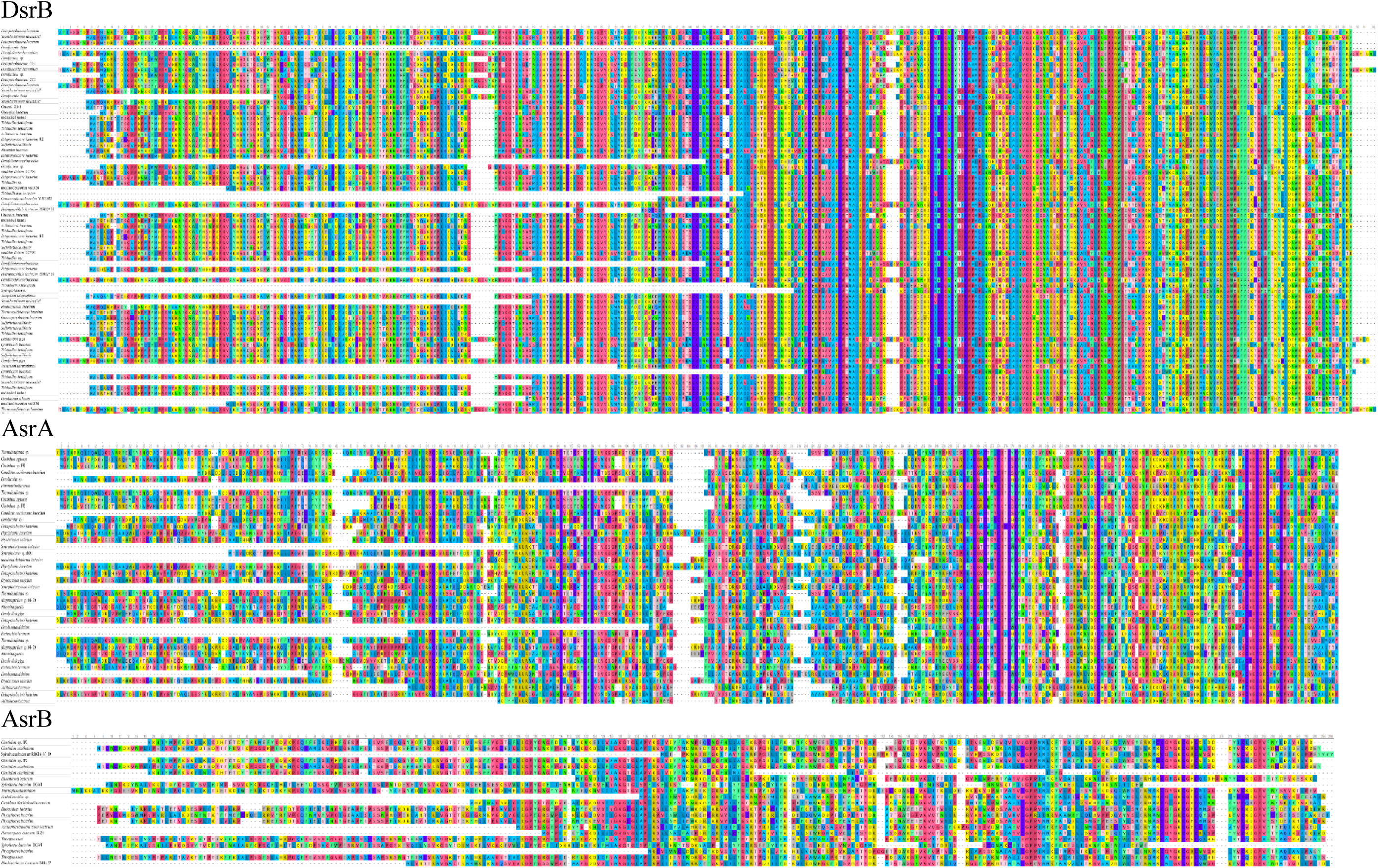
Sequence alignment of DsrA, DsrB, AsrA, AsrB proteins identified in this study. DsrA

**Supplementary File 8:**
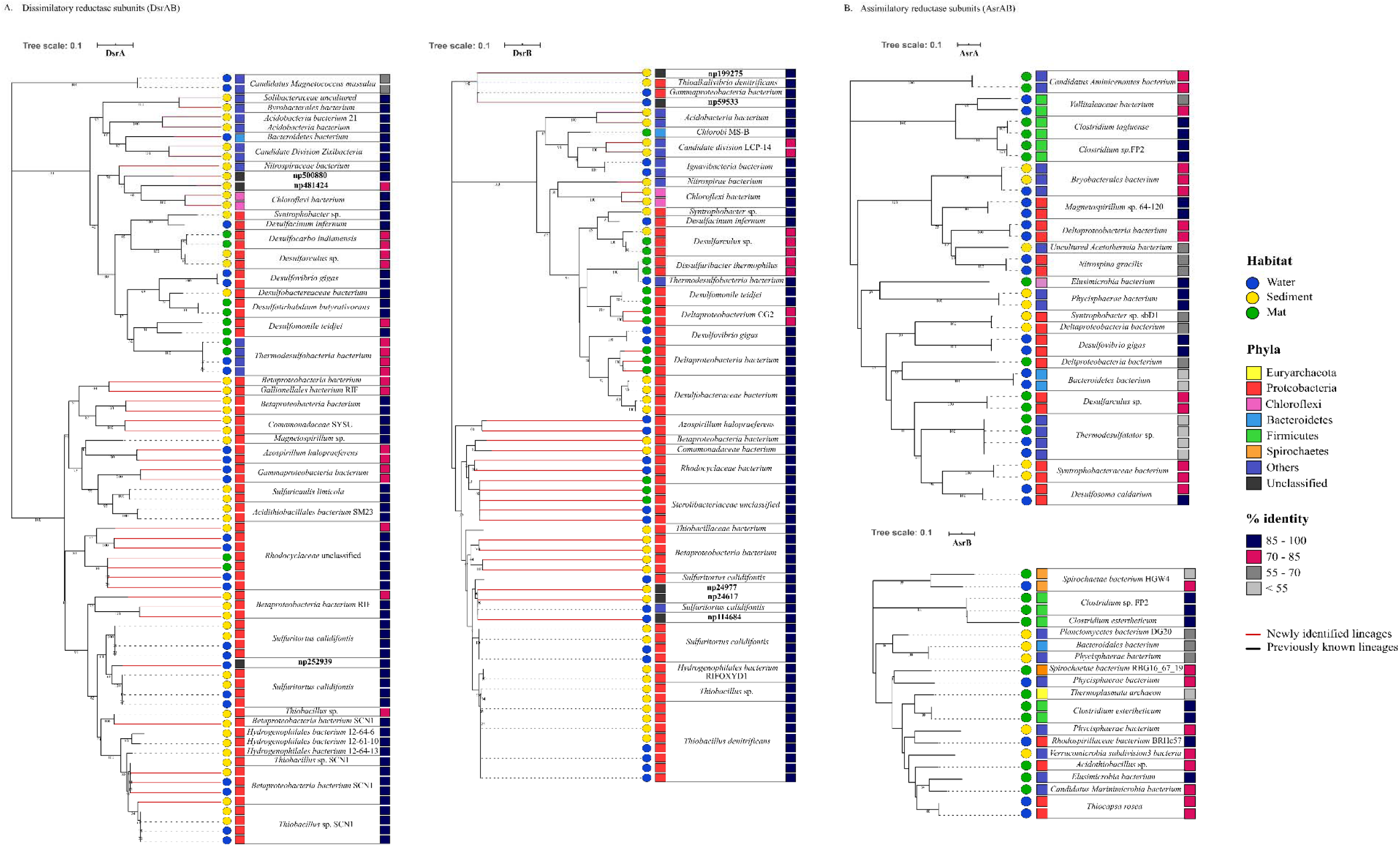
Phylogenetic analysis of DsrAB proteins: showing percentage similarity with taxa on ncbi *nr* database; Few newly identified lineages were marked in red. In DsrAB maximum no. of LGT events were occurred in Proteobacteria class itself *clade A2; B2* while the independent events of LGT towards different phyla was occurred through *A1, B1*. δ-Proteobacteria depicted in *clade*

**Supplementary File 9:**
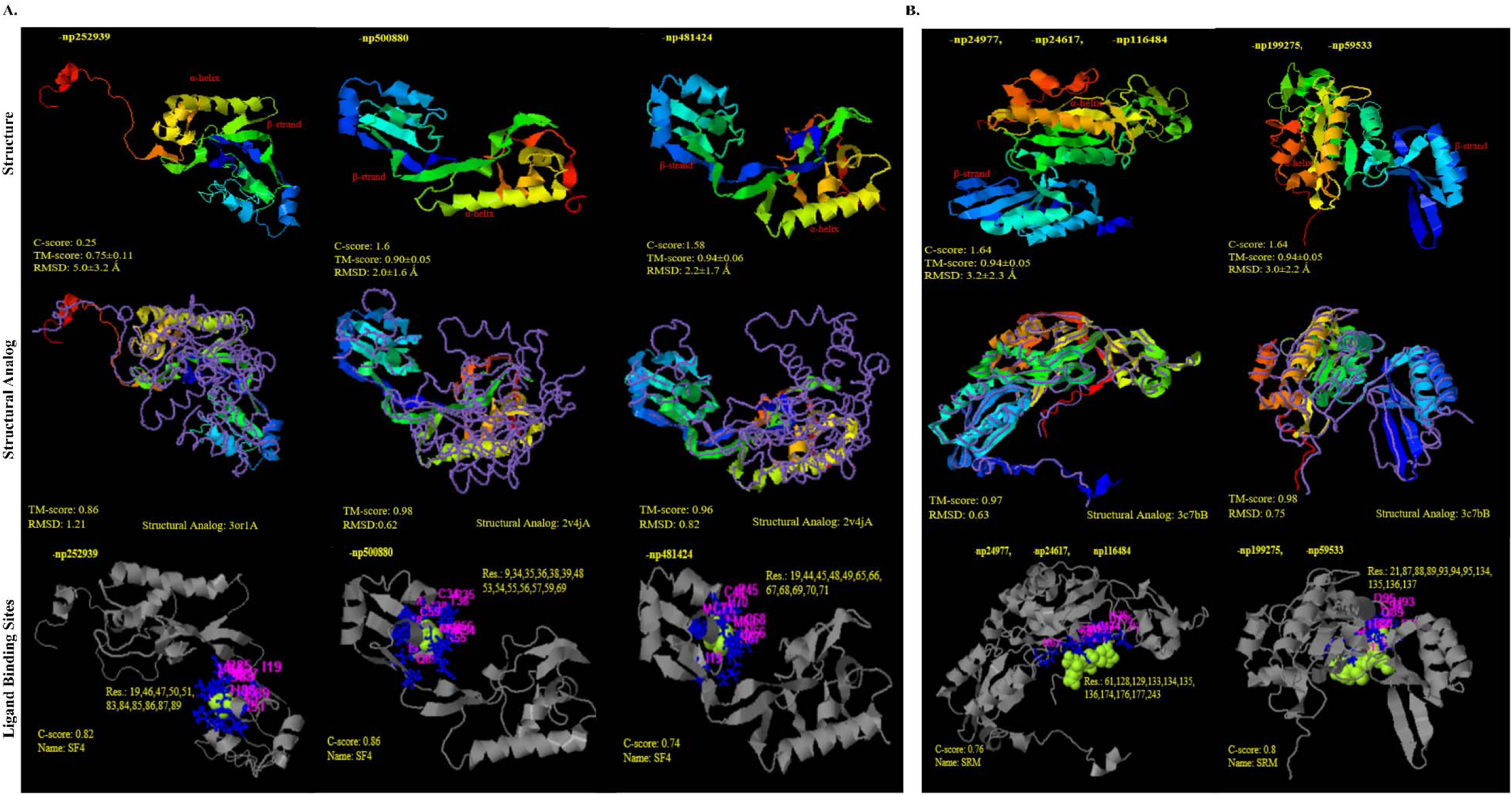
Structural similarity and analogy of unclassified proteins from PDB (Protein data bank) using i-TASSER. A) DsrA protein subunits i) np252939 and ii); iii) np500880; np481424 showing Tm-align similarity with PDB ids 3or1 and 2v4j (*Desulfovibrio gigas* and *Desulfovibrio vulgaris)* respectively; with SF4 (iron sulfur cluster) ligand binding sites B) DsrB protein subunits i) np24977, np24617, np116484 and ii) np199275, np59533 showing Tm-align similarity with PDB ids 3c7b; with SRM (siroheme) ligand binding sites.

**Supplementary File 10:**
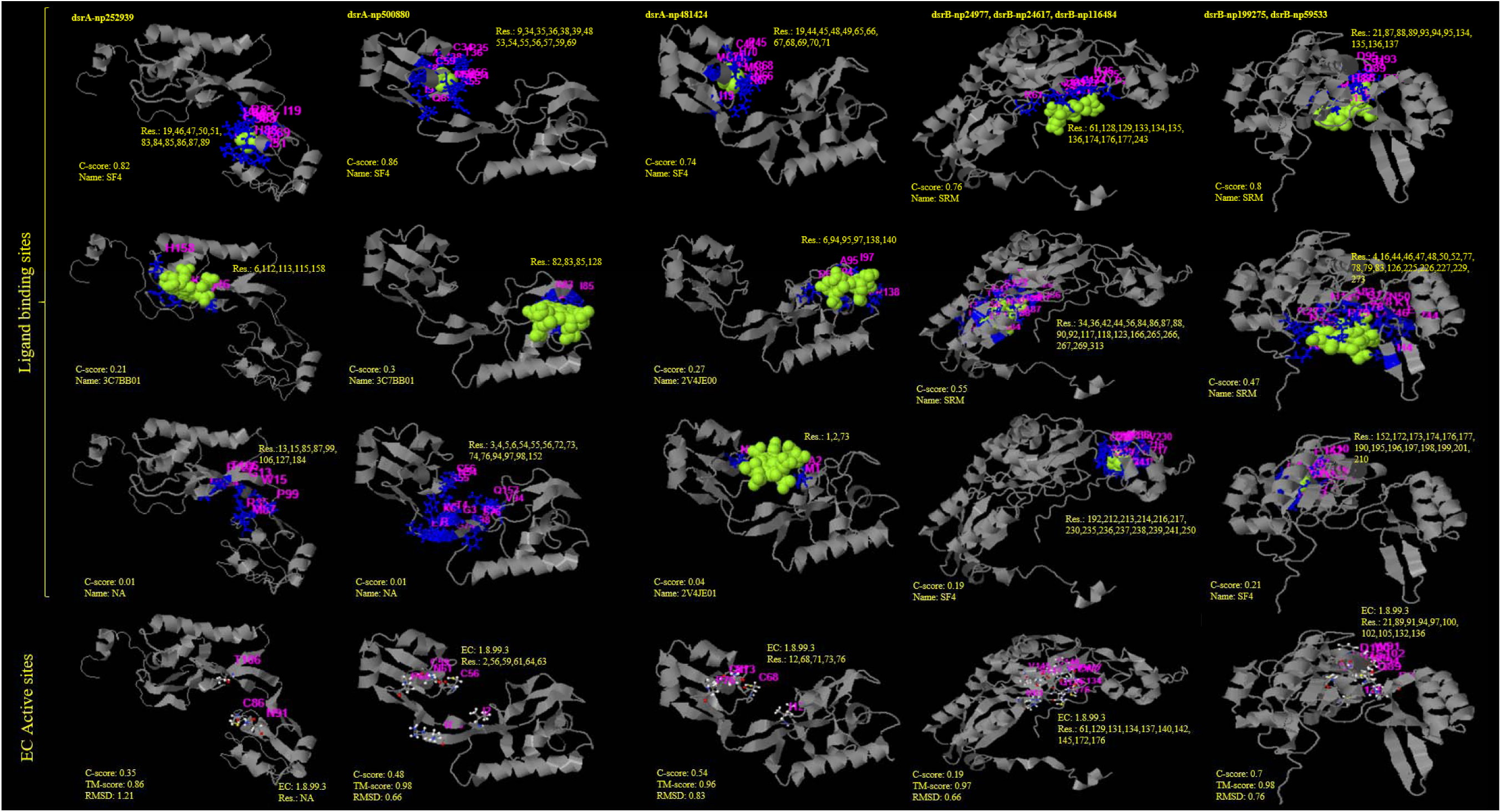
Structural models for other ligand binding sites of unclassified proteins of DsrA and DsrB subunits.

